# Form I and II Rubiscos Exhibit Temperature Dependent Carbon Kinetic Isotope Effects

**DOI:** 10.64898/2026.06.29.735352

**Authors:** Renée Z. Wang, Albert K. Liu, Patrick M. Shih, Daniel A. Stolper

## Abstract

Nearly all carbon on Earth today is fixed by the enzyme ribulose-1,5-bisphopshate carboxylase/oxygenase (‘rubisco’), which converts carbon dioxide (CO_2_) to sugar phosphates. All rubiscos measured thus far display a kinetic isotope effect (KIE) where ^12^CO_2_ is fixed at a faster rate than ^13^CO_2_. The relationship between rubisco’s KIE and the carbon isotope composition of plants, algae, and organic matter is central to many fields in the Earth sciences, plant biology, and biochemistry. Currently, all applications assume that the KIE does not vary with temperature. Here, we examine this assumption experimentally with *in vitro* KIE measurements of two rubiscos from phylogenetically distinct host organisms and rubisco protein clades – a Form I rubisco from the plant, *Spinacia oleracea* (spinach) and a Form II rubisco from the bacterium *Rhodosprillium rubrum*. We that find that both KIEs decrease linearly by ∼4.5‰ from 10-35°C with statistically indistinguishable slopes. We place these results into biological and geologic contexts by comparing them to observed variations in the carbon isotope composition of modern terrestrial plants and marine organic carbon, the geologic carbon isotope record, and rubisco’s biochemistry. We show that the measured temperature dependencies are sufficiently large to impact our interpretations of the enzymatic processes that drive variations in rubisco KIEs, as well as applications of stable carbon isotopes in the Earth and biological sciences.

**Significance Statement:** The carbon isotope composition of plants, algae, and organic matter are interpreted with models that assume the kinetic isotope effect of the carbon-fixing enzyme rubisco is temperature-independent, even though temperature varies by tens of degrees across the Earth today and in the past. Here, we demonstrate that the kinetic isotope effect of rubisco is temperature-dependent, suggesting that some of this isotopic variation may be due to intrinsic enzyme properties alone. In addition, though the rubiscos we measured are from diverse organisms (plant vs. bacteria), their KIEs show statistically indistinguishable temperature dependencies. This data forms the basis for future thermodynamic models on rubisco biochemistry.

## 1. Introduction

Nearly all carbon on Earth today is fixed by oxygenic photosynthesizers using the enzyme ribulose-1,5-bisphopshate carboxylase/oxygenase (‘rubisco,’ EC 4.1.1.39) (1), which initiates the Calvin cycle. Rubisco catalyzes both the carboxylation and oxygenation of the sugar phosphate ribulose-1,5-bisphosphate (RuBP) to yield two moles of 3-phosphoglyceric acid (3PGA) or one mole of 3PGA and one mole of 2-phosphoglycolate (2PG) respectively. Rubisco is not a singular enzyme, but rather an enzyme family composed of numerous clades (Fig. 1A). Only those from the Form I and II clades are known to be used in the canonical Calvin cycle to fix CO_2_ (2). Form I rubiscos are the most ecologically dominant rubisco clade on Earth today, used mainly by plants and algae as part of oxygenic photosynthesis (1, 3). In contrast, Form II rubiscos are primarily used by bacteria and archaea performing anoxygenic photosynthesis (though see (4) for exceptions).

**Figure 1:**
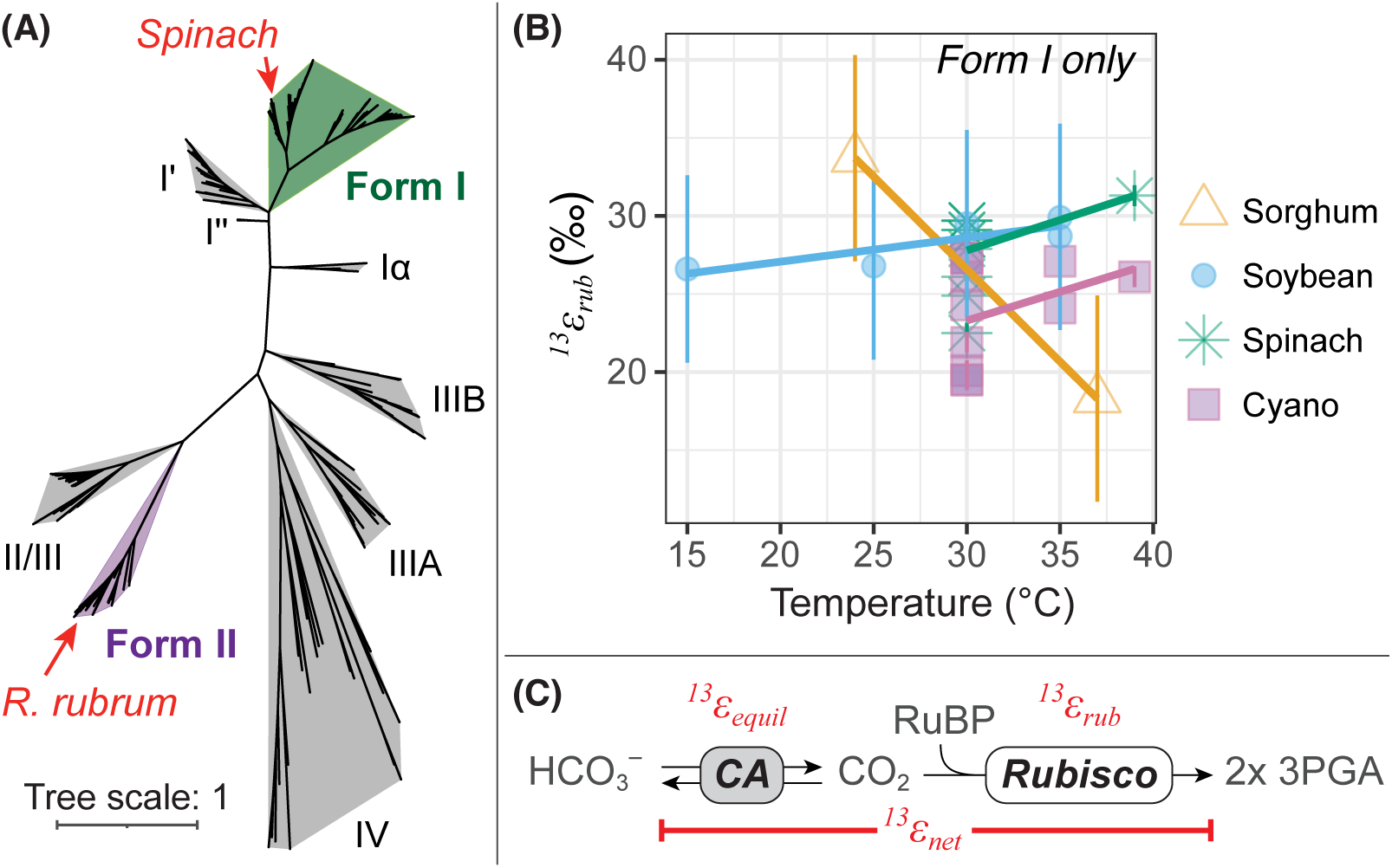
Temperature-dependence of carbon kinetic isotope effects in the rubisco superfamily is unclear. **(A)** Unrooted maximum likelihood phylogeny of rubisco’s large subunit sequence (*RbcL*, *n*=235) from (84). **(B)** Prior *in vitro* carbon KIEs (*^13^ε_rub_*) of Form I rubiscos where temperature was modified as a parameter: sorghum (31), soybean (33), spinach and the cyanobacterium *Synechococcus elongatus* PCC 7002 (32). Error bars (*s.d.*) are either reported in a given study (31) or calculated by us based on their reported data (32, 85). **(C)** Schematic representation of the processes that must be considered when calculating Rubisco ^13^KIE assays including the measured net fractionation (*^13^ε_net_*) and contribution of the equilibrium carbon isotope effect (*^13^ε_equil_*). CA: Carbonic anhydrase.

Every rubisco measured thus far exhibits a normal carbon kinetic isotope effect (^13^KIE) for carboxylation such that ^12^CO_2_ is converted into 3PGA at a faster rate than ^13^CO_2_ (*^12^k*>*^13^k*). These fractionations are reported in both alpha (*^13^α_rub_*) and epsilon (*^13^ε_rub_*) notation, where *^13^α_rub_* = *^12^k*/*^13^k* and *^13^ε_rub_* = (*^13^α_rub_* – 1)×1000 (5). In this convention, larger *^13^ε_rub_* values in units of per mil (‰) correspond to larger differences in the rate of ^12^C/^13^C uptake, with ^12^C always taken up faster than ^13^C. Rubisco’s ^13^KIE is generally thought to be the main source of the relative depletion of ^13^C observed in both modern and ancient organic carbon compared to inorganic carbon pools. The carbon isotope composition of these pools is reported in delta notation (δ) where δ^13^C = (*^13^R_sa_*/*^13^R_VPDB_* – 1) ×1000 and *^13^R* is the ^13^C/^12^C ratio in the sample relative to the VPDB reference scale. The difference in carbon isotope composition between dissolved CO_2_ (‘*d’*) and the primary photosynthate (‘*p*’) is defined as *^13^ε_P_* ≡ (*^13^R_d_*/*^13^R_p_* – 1) ×1000 ≍ δ^13^C_d_ –δ^13^C_p_ (6); the carbon isotope composition of organic carbon (δ^13^C_org_) is typically depleted by 10s of per mil relative to source inorganic carbon.

The linkage between δ^13^C_org_ and rubisco’s ^13^KIE forms the basis of many applications of stable carbon isotopes in both the Earth and biological sciences. For example, in the Earth sciences, this linkage has been used to reconstruct concentrations of atmospheric oxygen (e.g., (7)) and carbon dioxide (8) over geologic time, identify the earliest evidence of life on Earth (e.g., (9, 10)), and reconstruct the diets of ancient organisms (e.g., (11)). In biology, these isotope effects are commonly to distinguish C3 vs. C4 plant metabolisms (e.g., (12)) and estimate plant water use efficiency and internal plant *p*CO_2_ concentrations (e.g., (13)). In biochemistry, rubisco’s ^13^KIE has been used as a constraint on rubisco’s reaction mechanism (14–18). However, despite the importance of rubisco’s ^13^KIE in diverse fields, our understanding of the biochemical, phylogenetic, and environmental controls that set the variation and magnitude of rubisco’s ^13^KIE has only been explored across a limited diversity of organisms and in an even more limited range of experimental conditions (see (19–21) for recent compilations).

Rubisco ^13^KIEs have been observed to vary by ∼20‰ (∼10-30‰) across the 17 wild-type (WT) host organisms examined thus far, 14 of which are from the Form I clade (Supplemental Dataset 1). It is generally thought that this variation arises from biochemical differences among rubiscos derived from phylogenetically distinct clades (14–18). The value of rubisco’s ^13^KIE from a given organism measured *in vitro* is generally assumed (often implicitly) to be representative under all conditions in natural environments, even though these experiments are always conducted under similar conditions at room temperature (∼20°C), elevated dissolved inorganic and Mg^2+^ concentrations (order 10 mM), and neutral to slightly alkaline pH.

The question we take up here is whether these experimental parameters influence the measured ^13^KIE with a focus on temperature-dependent effects. We are motived to examine the role of temperature for two reasons: First, the catalytic activity of rubisco, like all enzymes, increases with increasing temperature for both carboxylation and oxygenation until reaching a peak prior to denaturing (22). Second, the temperatures at which oxygenic photosynthesis takes place vary by ∼20-30°C across the Earth today ((23, 24), see Discussion for more detail) and have been reconstructed to have varied significantly in the past. For example, mean sea surface temperatures are thought to have varied between 10°C to 35°C over the past ∼500 million (25) and some reconstructions place average Archean and Proterozoic ocean temperatures above 50°C (540-4000 million years ago; (26–28)), though we note much disagreement on this topic exists (i.e., (29, 30)). As such, if rubisco exhibits a temperature dependent ^13^KIE, it could influence our understanding of rubisco’s biochemistry as well quantitative interpretations of modern and ancient δ^13^C_org_.

Whether or not rubisco exhibits a temperature dependent ^13^KIE was last studied in a series of papers from 1960s-1980s. These were conducted on Form I rubiscos and disagreed both in the magnitude and sign of the temperature dependence (-1.18 to 0.28 ‰/°C; Fig. 1B (31–33)). This disagreement has generally been attributed to artifacts associated with the experimental techniques used at the time (13, 34, 35) and the results from these experiments are typically not considered. Consequently, the foundational mathematical models that directly relate rubisco’s ^13^KIE to the δ^13^C of plants (e.g., (13)) assume that rubisco’s ^13^KIE is temperature independent. This assumption of temperature independence has persisted in effectively all subsequent models and interpretations rubisco’s influence on the δ^13^C of photosynthate and biomass in not only plants, but also aquatic microorganisms such as cyanobacteria and algae (e.g., (36, 37)).

Here, we directly test whether rubisco does or does not express a temperature dependent ^13^KIE. Specifically, we measured the temperature dependence of *^13^ε_rub_* from 10- 35°C of using two common but distantly related rubiscos, that from *Spinacia oleracea* (“spinach,” a model Form I) and *Rhodospirillum rubrum* (a model Form II). Our results demonstrate that a clear and significant temperature dependence exists for both phylogenetically distinct forms of rubisco.

## 2. Results

We determined values of *^13^ε_rub_* following the “substrate depletion method” (e.g., (38, 39)) in which the δ^13^C of dissolved inorganic carbon (DIC) is measured as a function of its fractional uptake and then fit to a Rayleigh model. Specifically, we regressed the natural logarithm of ^13^C/^12^C (*^13^R*) of DIC normalized to its initial composition vs. the fraction of DIC remaining (*f*). The slope of this line corresponds to the net KIE (*^13^ε_net_*) associated with CO_2_ uptake and fixation by rubisco. Here, *^13^ε_net_* includes contributions from both rubisco (*^13^ε_rub_*) and the known temperature-dependent equilibrium isotope effect between HCO_3_^-^ (the dominant DIC species at pH 8; 98%) and CO_2(aq)_ (*^13^ε_equil_*; (40), Fig. 1C). We correct *^13^ε_net_* with *^13^ε_equil_* to find *^13^ε_rub_* (Methods).

Our replicate experimental determinations of *^13^ε_rub_* for spinach at room temperature and 35°C and *R. rubrum* at 35°C yielded a pooled standard deviation of 1.0‰, which we take as an estimate for our complete procedural reproducibility. We verified accuracy by comparing our measured *^13^ε_rub_* values to prior determinations at room temperature (i.e., ∼25°C). Our replicate values of *^13^ε_rub_* at room temperature for spinach are 26.8±0.7‰ (1SE) and 27.4±0.8‰ (Fig. 2A, Table S1), which agree with prior determinations of 26.4 to 30.4‰ (Fig. 2B, Dataset S1, (19, 38, 41–43)). For *R. rubrum*, we determined a room temperature value of 19.2±1.0‰ (1SE), which again falls within the determined range of 17.8‰ to 23.0‰ (Fig. 2B, Dataset S1, (38, 44, 45)). Based on this, we proceed with the assumption that our methodology yields accurate results in so far as that they are indistinguishable from prior measured ranges of *^13^ε_rub_* from the same organisms.

**Figure 2:**
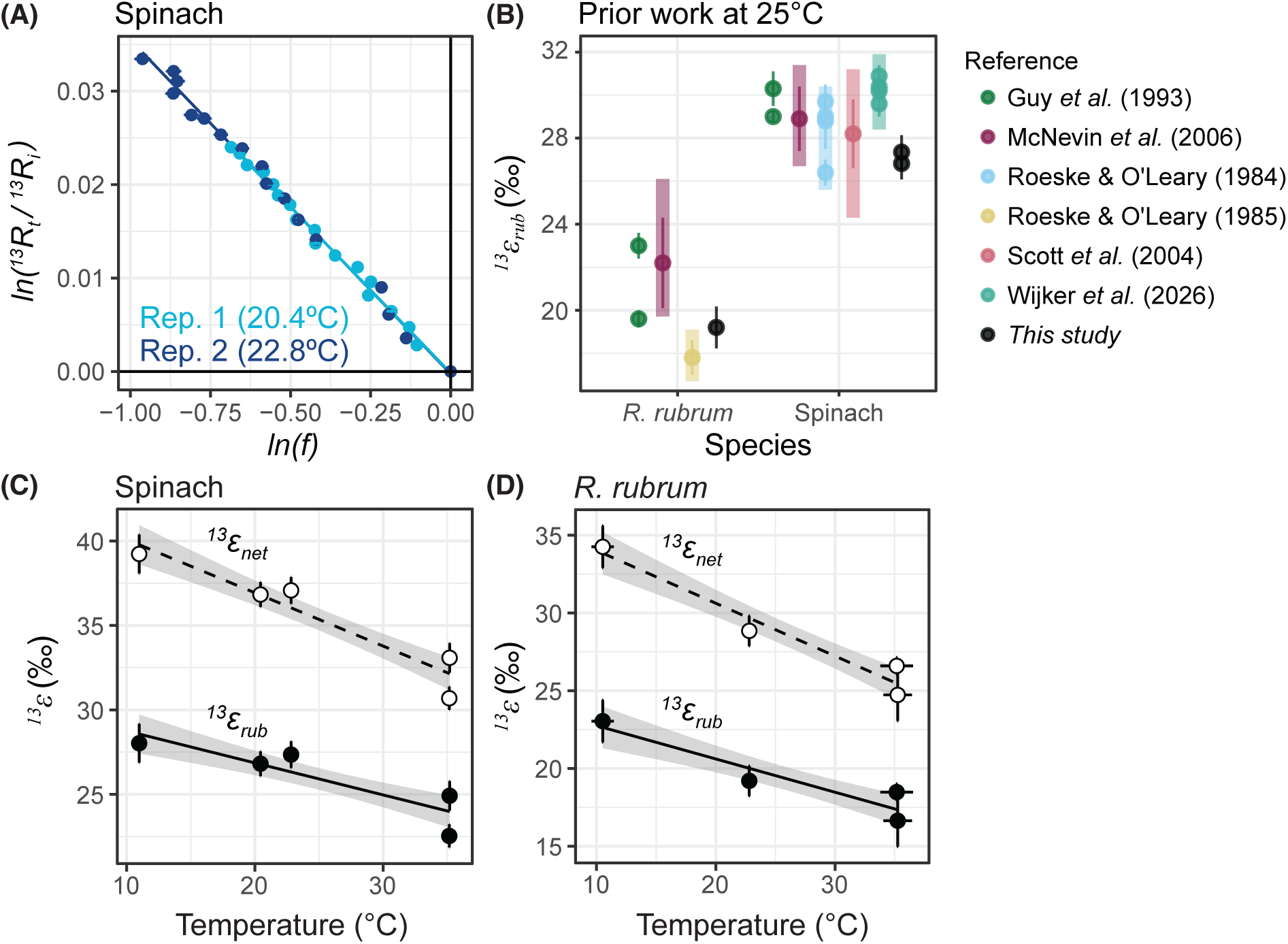
Results. **(A)** Rayleigh curves of replicate spinach experiments at room temperature from this study; linear regression is shown without uncertainty for figure clarity. Error on individual points indicates ±1SE. **(B)** Comparison of room temperature (20-23°C) *R. rubrum* and spinach measurements to previously published *in vitro* experiments performed at room temperature (19, 38, 41, 43–45). **(C,D)** Net fractionation (*^13^ε_net_*, open circles, dashed line) and rubisco fractionation (*^13^ε_rub_*, closed circles, solid line) for spinach and *R. rubrum* respectively. Error bars on data points indicate ±1SE uncertainty and error envelope on linear regression shows 68% c.i. See Table 1 for linear regression results.

**Table 1:** Linear regressions for *^13^*ε vs. temperature (°C). Best fits for *^13^*ε = *m*\*T + *b*, where temperature is in Celsius. Reported uncertainty on *m* and *b* is ± 1SE from a simple linear regression. The covariance on the slope and intercept (σ*_mb_*) is -0.84. Linear regression and calculation of covariance matrix was performed in RStudio (v2023.06.1+524) with R (v4.4.0) [call: *lm()*; *vcov()*] (90).

| Strain | Fractionation | Slope ($m$) | | Intercept ($b$) | | Adj. $R^2$ |
| --- | --- | --- | --- | --- | --- | --- |
| | | Estimate | $\pm 1\text{SE}$ | Estimate | $\pm 1\text{SE}$ | |
| Spinach | Net ($^{13}\epsilon_{net}$) | -0.32 | 0.06 | 43.25 | 1.55 | 0.876 |
| | Rubisco ($^{13}\epsilon_{rub}$) | -0.19 | 0.06 | 30.65 | 1.54 | 0.707 |
| *R. rubrum* | Net ($^{13}\epsilon_{net}$) | -0.34 | 0.06 | 37.42 | 1.56 | 0.923 |
| | Rubisco ($^{13}\epsilon_{rub}$) | -0.21 | 0.06 | 24.89 | 1.54 | 0.824 |

In terms of temperature dependence, we find that for both spinach and *R. rubrum*, *^13^ε_rub_* decreases with increasing temperature (Fig. 2C,D; Table 1). Specifically, spinach’s *^13^ε_rub_* decreases from 28.0‰ at 10°C to 23.7‰ at 35°C (-0.19/°C) while *R. rubrum*’s *^13^ε_rub_* decreases from 23.0‰ at 10°C to 17.6‰ at 35°C (-0.21/°C). These slopes are statistically identical both as ‰/°C (*P*=0.77) or as *ln(^13^α_rub_)* vs. 1/T (T in Kelvin; *P*=0.58; Fig. S1). As such, the ∼8‰ difference in *^13^ε_rub_* between spinach and *R. rubrum* at room temperature is preserved across the temperature range examined. In addition, the sign of the temperature dependence of *^13^ε_rub_* is the same as that for *^13^ε_equil_* such that the temperature dependence of *^13^ε_net_* is larger than either of the two separately. Specifically, *^13^ε_net_* decreases by -0.32/°C for spinach and -0.34/°C for *R. rubrum*.

## 3. Discussion

As introduced above, prior work has assumed that any variations in *^13^ε_p_* (i.e., the difference in δ^13^C between organic carbon and substrate) in space or time represent physiological responses to changes in the rate limiting step of carbon fixation (e.g., diffusion vs. carbon fixation by rubisco). Such can occur, for example, due to changes in water availability or CO_2_ concentrations (e.g., (13, 37)). Embedded in this interpretation of *^13^ε_p_* is that, for a given organism, the intrinsic ^13^KIE of rubisco *itself* does not vary. Our experiments indicate that this assumption does not hold for rubisco derived from two distinct species and enzyme clades (Form I and Form II; Fig. 1A). These yield measurable changes in ^13^KIEs from 10 to 35°C that correspond to a decrease of ∼5‰ for *^13^ε_rub_* and ∼8-9‰ for *^13^ε_net_*. As we will now discuss, these changes are large enough that they have implications for interpretations of terrestrial and marine δ^13^C_org_ today and in the geologic past. We structure our discussion by first examining the implications for modern terrestrial and marine systems, then moving to the geologic past, and finally closing with an examination of the work in the context of rubisco’s biochemistry.

### 3.1 Modern Terrestrial Systems

We examine the implications of our results for terrestrial systems by quantifying potential changes in the δ^13^C of plant photosynthate due solely to the temperature dependence of *^13^ε_rub_*. To do this, we estimate the range of mean plant growing season temperatures (*T_gs_*) using the global leaf data set of (24) as a proxy for average photosynthesis temperatures (Fig. 3A, Methods). We calculate *^13^ε_rub_* as a function of *T_gs_* (Fig. 3B) using our measurement of spinach rubisco, which is the model for C3 plant ^13^KIEs. In doing this, we do not include contributions of equilibrium DIC fractionations (i.e., *^13^ε_equil_* to calculate *^13^ε_net_*), because the source of inorganic carbon for leaves is atmospheric CO_2_.

**Figure 3:**
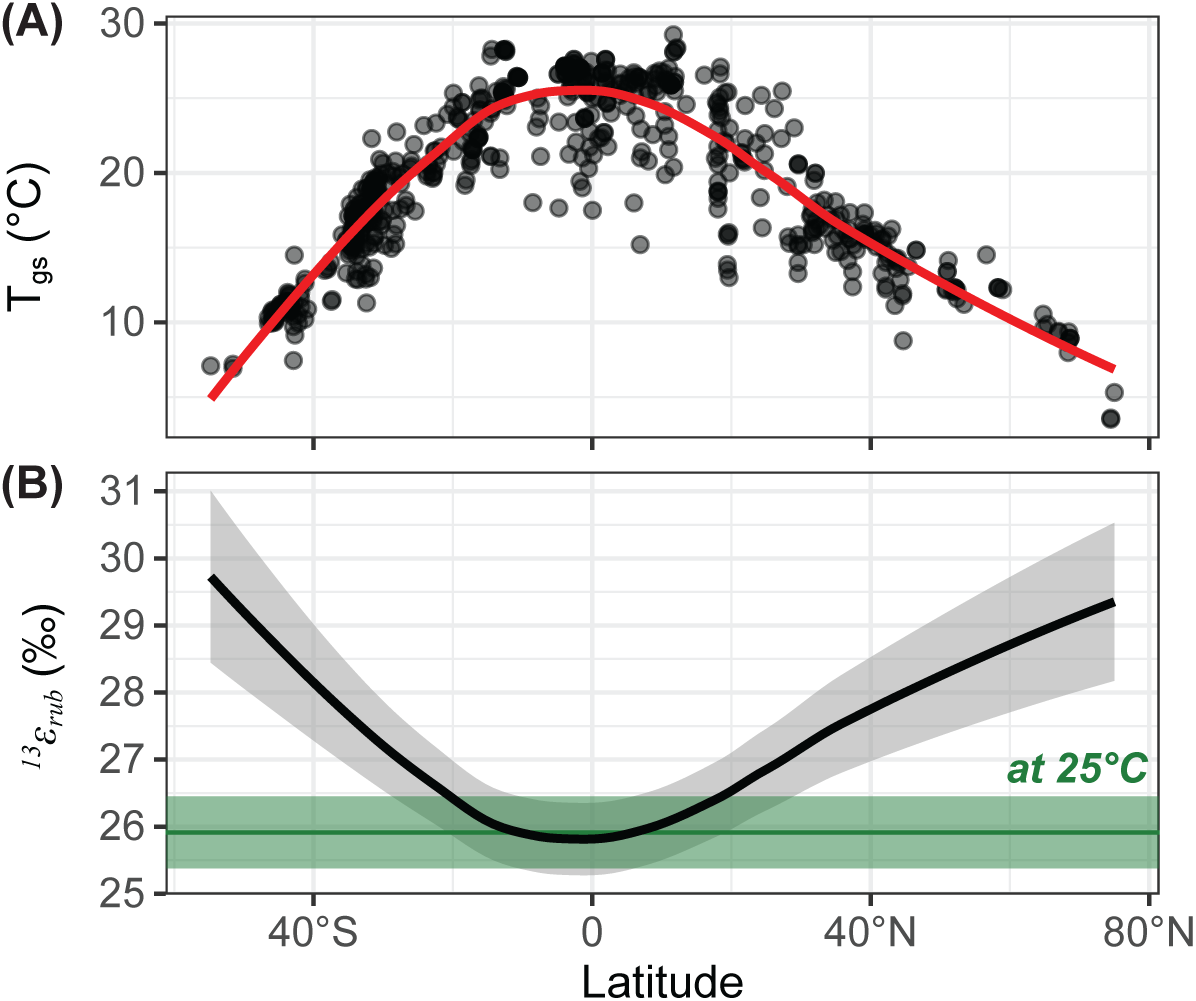
Predicted carbon kinetic isotope effects values for spinach across latitudes. **(A)** Mean growing season temperatures (*T_gs_,* black circles, *n*=655) from (24) for elevations <2,000 m to restrict the effects of elevation on temperature). Data are fit to a smoothed curve (“LOESS”; red line). **(B)** Spinach *^13^ε_rub_* (mean ±1SE) for LOESS-smoothed mean *T_gs_*, assuming no uncertainty on temperature and using the temperature-dependence measured in this study. Green line and shaded envelope (±1SE) indicate value at 25°C, calculated from our data.

We find that *^13^ε_rub_* ranges from 26-30‰ from the equator to high latitudes, which corresponds with variations in *T_gs_* from 5-25°C (Fig. 3B). This ∼4‰ range is of similar order to global variations in bulk plant δ^13^C (46–48). The full isotope effect of rubisco is not expressed by plants due to partial limitation of fixation by CO_2_ diffusion. To take this into account, we took our results and placed them into a standard model of plant carbon isotope fractionation (13) with typical ratios of internal to external CO_2_ concentrations (∼0.6 to 0.7, (13, 49)). Including this decreases the total variation in plant δ^13^C_org_ that may be attributed to the ^13^KIE’s temperature dependence to ∼2-3‰.

Variations in plant δ^13^C_org_ across the globe are largely attributed to variations in water availability via its effects on stomatal processes (46–48). More recently, it has also been proposed that leaf δ^13^C is also modulated by fluxes of isoprene out of the leaf that increase with temperature; this would then cause leaf δ^13^C to increase with temperature (opposite temperature trend of *^13^ε_rub_* measured here; (50)). Our measured temperature dependence of *^13^ε_rub_* does not run counter to any of these proposals, and they are well supported by observations and models. Rather, our results indicate that an additional parameter should also be considered when evaluating the processes that set the δ^13^C_org_ of plants across Earth’s terrestrial ecosystem. The effects of temperature can be explored straightforwardly by modifying existing isotopically enabled models of terrestrial carbon fixation to include the temperature-dependence rubisco’s ^13^KIE. Such models will be needed to evaluate when temperature vs. other parameters dominate any observed changes in δ^13^C_org_ in terrestrial ecosystems.

### 3.2 Modern Marine Systems

We now turn to modern marine systems and examine the implications of our experiments in a similar manner as done for terrestrial systems above. To do this, we first estimated the range of temperatures at which marine photosynthesis occurs today. Specifically, we weighted sea-surface temperatures (SSTs) with monthly estimates of net primary productivity (NPP) taken from the Carbon-based Production Model from (51) (Methods; Fig. S2). We use NPP instead of gross primary productivity as it integrates both carbon fixation and export out of the photic zone and thus captures the organic carbon that can eventually be preserved in the rock record. We find that the weighted SST of photosynthesis (SST_w_) varies from -1 to 28°C as a function of latitude (Fig. 4A).

**Figure 4:**
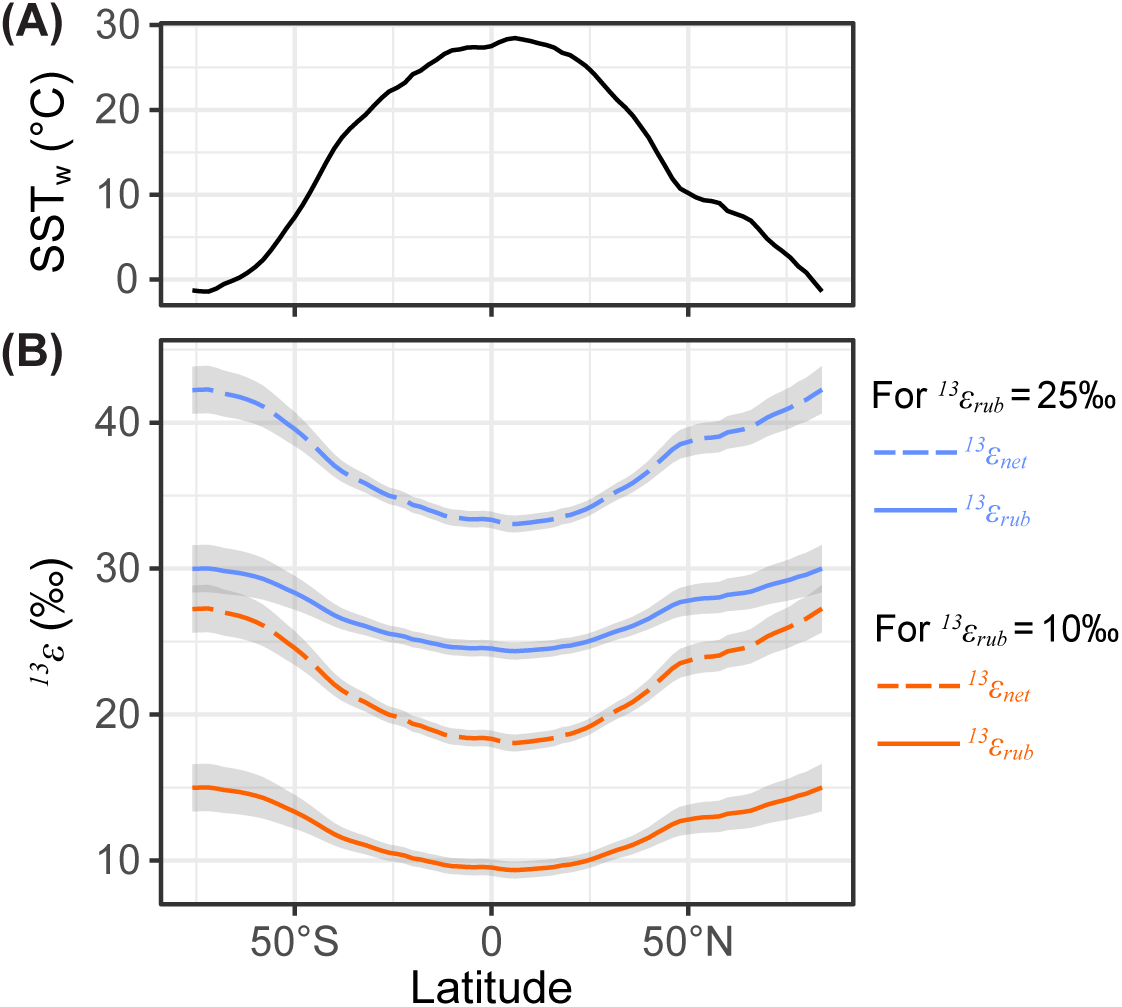
Predicted carbon kinetic isotope effects values for marine systems. **(A)** Sea-surface temperatures weighted by net primary productivity (SST_w_) for 2013-2022. **(B)** Solid lines with shaded envelope (±1SE) indicate latitudinal variation in *^13^ε_rub_* assuming *^13^ε_rub_* = 10‰ (blue) or *^13^ε_rub_* = 25‰ (orange) at 25°C, using the temperature-dependence of spinach and SST_w_ from (A). Dashed lines with shaded envelope (±1SE) show *^13^ε_net_*, which includes both the contributions of both rubisco and isotopic equilibrium between HCO_3_^-^ and CO_2(aq)_.

We next calculated the variation in *^13^ε_rub_* across this temperature range. As we did not measure any rubiscos from marine organisms in this study, we took the full range of marine *^13^ε_rub_* at 25°C (∼10-25‰, Supplemental Dataset S1) and assumed they have same temperature dependence as spinach. We consider this an acceptable assumption for two reasons – first, all ecologically dominant marine photoautotrophs (i.e., diatoms, coccolithophores) use Form I rubiscos, like spinach measured here. Second, the temperature dependence of *^13^ε_rub_* is statistically identical for Form I and Form II rubiscos even though they are phylogenetically distinct, exhibit unique oligomeric structures, and have different *^13^ε_rub_* values at 25°C. The latter predicts broadly similar temperature dependences for rubisco ^13^KIEs. However, this proposal will require additional experimental confirmation.

Given these assumptions, we find a total variation in *^13^ε_rub_* of 6‰ that is solely due to temperature. The absolute values range from 9-15‰ if one assumes *^13^ε_rub_* = 10‰ at 25°C, or 24-30‰ if one assumes *^13^ε_rub_* = 25‰ at 25°C (Fig. 4B,C). The net fractionation (*^13^ε_net_*), which includes the additional temperature-dependent isotope effects of the DIC system (*^13^ε_equil_*), varies by 9‰ over this temperature range with absolute values of 18-27‰ (*^13^ε_rub_* = 10‰ at 25°C) or 33-42‰ (*^13^ε_rub_* = 25‰ at 25°C). We consider *^13^ε_net_* the more representative parameter for interpreting marine δ^13^C_org_ as the isotopic composition of bicarbonate buffers the δ^13^C of DIC in marine systems such that the isotopic composition of CO_2_ can itself vary as a function of temperature.

Importantly, the results of our simplified calculation are consistent with the range of values of *^13^ε_p_* (on the order of 10‰; (52, 53)) and δ^13^C_POC_ (roughly 10‰; Fig. S3; (54–56)) observed in the surface oceans as a function of temperature. They also have the same sign (larger *^13^ε_rub_* for more depleted δ^13^C_POC_). Additionally, recent studies have found strong correlations between temperature and marine particulate organic carbon δ^13^C (57) and algal δ^13^C (58). We note there are other parameters that must also be considered when interpreting marine δ^13^C_org_, including latitudinal variations in the distribution of primary producers (each with their own distinct rubisco and corresponding ^13^KIEs), as well as the role of light intensity and CO_2_ concentrating mechanisms (CCMs) in potentially setting the δ^13^C of algae (e.g., (59–61)). Regardless, our results indicate that changes in *^13^ε_rub_* intrinsic to the enzyme, along with the equilibrium isotope effects in DIC, could be responsible for a significant portion of the variation in marine δ^13^C_org_ as a function of latitude.

### 3.3 The Geologic Record

We next examine the implications of our results for interpreting the geologic record of δ^13^C_org_. This record is central to the development of two important hypotheses about rubisco’s isotopic history. First, the depletions in δ^13^C_org_ of ancient vs. inorganic carbon are similar in magnitude to those observed on modern Earth, and these depletions and can be found in rocks at least as old as 3.8 billion years old (9) and even in graphitic inclusions in a 4.1-billion-year-old Hadean zircon (10). This similarity to the modern has led to the proposal that the Calvin cycle was not only present but was also the dominant carbon fixation pathway over the past 3.8 billion years. Such proposals require rubisco was the isotopically discriminating biosynthetic step of carbon fixation (62). Second, it has been proposed that rubisco’s ^13^KIEs has been approximately constant over billions of years (63) such that *^13^ε_p_* and *^13^ε_rub_* measured in modern organisms can be used directly to interpret the past. Implicit in both hypotheses is that *^13^ε_rub_* did not change over time despite potential large-scale changes in temperature over geologic history. Given our results, we now consider whether changes in temperature over geologic time are sufficient to influence interpretations of the geologic record of δ^13^C_org_.

To explore this, we calculate values of *^13^ε_rub_* and *^13^ε_p_* as a function of temperature and compared these to the geologic record of δ^13^C_org_ To do this, we made the following assumptions: First, we assumed the temperature dependence of spinach can be used to calculate *^13^ε_rub_* over geologic time with a fixed value of *^13^ε_rub_* = 25‰ at 25°C. This is a typical value assumed in geologic studies (25-28‰; e.g., (64)) and is similar to the average value for cyanobacterial rubiscos (median *^13^ε_rub_* of 25.3‰; *n* = 6; Supplementary Dataset S1), which are thought to have been the dominant primary producers for most of the Proterozoic (65). Second, we assumed that *^13^ε_rub_* is maximally expressed so that it is equal to *^13^ε_p_*. In doing this, we are effectively assuming that transport of inorganic carbon into the cell does not influence *^13^ε_p_*. Third, we assumed that HCO_3_^-^was always the dominant DIC species in marine waters. This assumption is consistent with prior estimates of seawater pH (6.5-8) over geologic time (66, 67). Fourth, we assume that the δ^13^C of the DIC pool (δ^13^C_DIC_) has been ∼0‰ over geologic time, which is consistent with compilations of carbonate δ^13^C over geologic time ((68, 69); Fig. 5). As such, this analysis does not consider spatial variations in δ^13^C or times when carbonate δ^13^C changed rapidly (e.g., various carbon isotope excursion). The third and fourth assumptions allow us to calculate the δ^13^C of CO_2(aq)_ at a given temperature relative to seawater HCO_3_^-^ with δ^13^C of 0‰.

**Figure 5:**
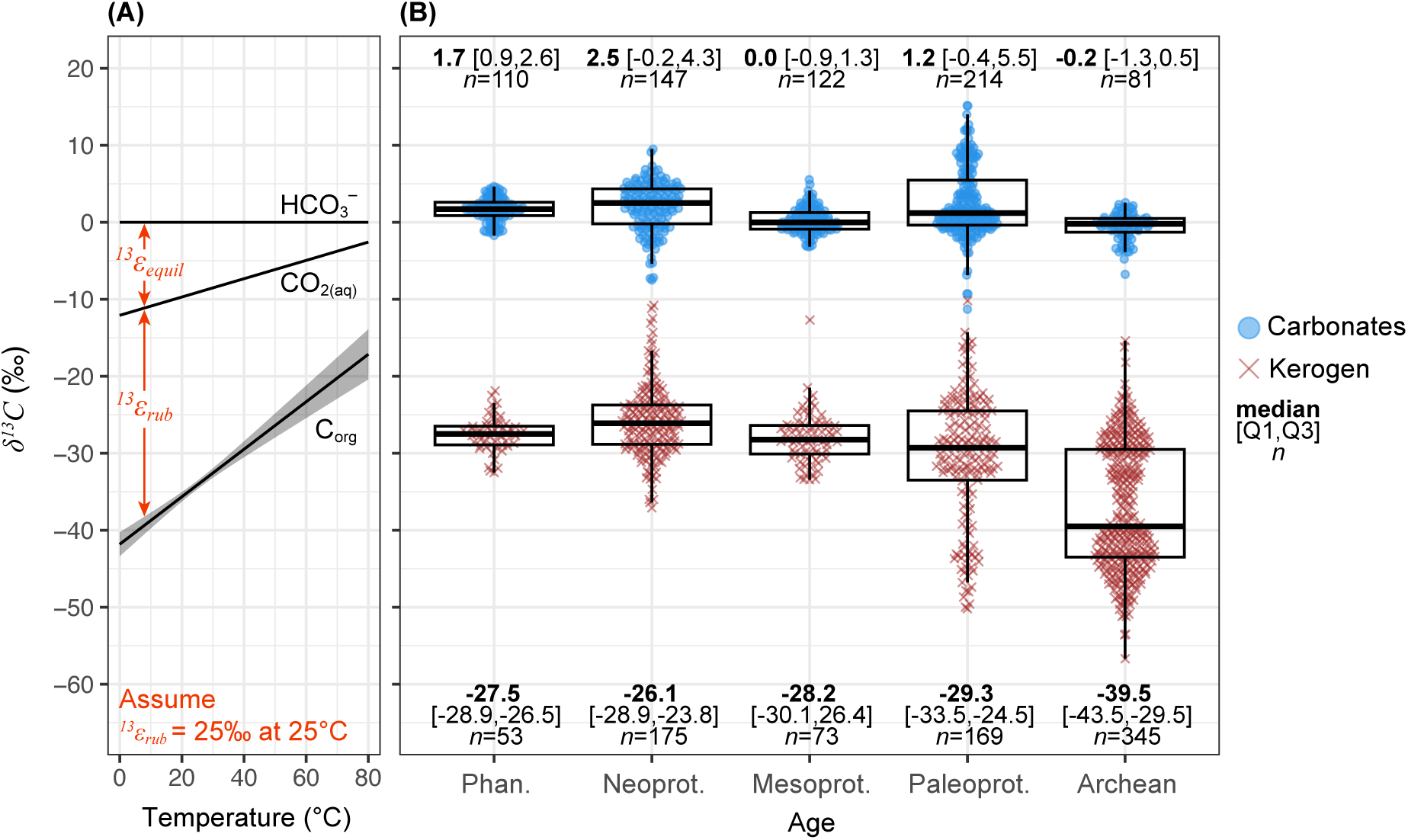
Isotope composition of organic carbon over geologic time. **(A)** Carbon isotope composition (δ^13^C) of bicarbonate (HCO_3_^-^), aqueous CO_2_ (CO_2(aq)_) and organic carbon (C_org_) as a function of temperature, assuming *^13^ε_rub_* = 25‰ at 25°C. The temperature-dependence of *^13^ε_rub_* is taken from spinach determined here and the equilibrium isotope effect (*^13^ε_equil_*) from (40). **(B)** Carbonate (blue circles) and kerogen data (brown crosses) are taken from the compilation in (69). Carbonate data is restricted to samples labeled as carbonates, limestones, and/or dolomites, while kerogen data is restricted to metamorphic grades of greenschist and below. Tukey boxplots (black) show the median and interquartile range (IQR) from 25^th^ to 75^th^ percentile (Q1,Q3); whiskers indicate 1.5 times the IQR from the quartiles. Statistics are given as: median [Q1,Q3] *n* (sample number). Geologic ages abbreviated as Phan. (Phanerozoic), Neoprot. (Neoproterozoic), Mesoprot. (Mesoproterozoic), Paleoprot. (Paleoproterozoic).

With these assumptions in mind, we calculate δ^13^C_org_ from 0-80°C and compare them to compiled values of kerogen δ^13^C taken from (69) (δ^13^C_kerogen_; Fig. 5). A temperature range of 0-80°C was used as this encompasses the modern SST range (∼0-25°C), the range of average SSTs over the Phanerozoic (∼10-35°C; e.g., (25)) and typical maximum estimate of SSTs for the Archean (e.g., (26)), though we again stress there is not a consensus on Precambrian SSTs. Compiled δ^13^C_kerogen_ values are taken only from greenschist or lower metamorphic facies to avoid metamorphic effects on δ^13^C.

We calculate that δ^13^C_org_ ranges from -42 to -17‰ from 0-80°C, with lower δ^13^C values at lower temperatures (Fig. 5A). Over the Phanerozoic, compiled δ^13^C_kerogen_ data have a median value of - 27.5‰ (Fig. 5B), which corresponds to ∼46°C based on our assumptions. This temperature is 10°C above the 11-36°C range of recent Phanerozoic temperature reconstructions (25) and far outside of any predicted mean temperature of the Earth over this time frame. If CO_2_ is not limiting (i.e., *^13^ε_rub_* sets *^13^ε_p_*), then the global average value of *^13^ε_rub_* would need to be <22‰ at 25°C to fall within a 11-36°C temperature range. Though possible, an alternative interpretation is that *^13^ε_rub_* is not fully expressed during carbon fixation, either due to isotope effects associated directly with CO_2_ limitation or indirectly via the effect of CCMs (36, 37, 59–61).

The median δ^13^C_kerogen_ of the Proterozoic Eras range from -26.1 to -29‰ (Fig. 5B). This corresponds to mean temperatures of 40-50°C. This would be consistent with some reconstructions of a warm Proterozoic climates (26–28). Alternately, as in our interpretations of the Phanerozoic, the ^13^KIE of rubisco could have been smaller (δ^13^C_org_ = -28.2‰ if *^13^ε_rub_* = 20‰ at 25°C), rubisco was limited by CO_2_ availability, and/or CCMs had evolved and were already significantly impacting marine δ^13^C_org_ in the Proterozoic.

Finally, the median value of δ^13^C_kerogen_ in the Archean is -39.5‰ (Fig. 5B). The distribution is bimodal with peaks around -28 and -44‰. Why Archean δ^13^C_kerogen_ is typically lower than Proterozoic and Phanerozoic equivalents is not fully understood. Our model returns a temperature of ∼8°C for the median δ^13^C_kerogen_ of -39.5‰. The temperatures are 45°C and <0°C for the two bimodal peaks. As such, the simplified framework used here would generally require that ancient organisms grew at temperatures <50°C, which is inconsistent with some of the highest estimates of Archean SSTs (e.g., ∼80°C). Alternatively, Archean rubiscos could have had significantly larger values of *^13^ε_rub_* and/or temperature-dependencies than those used in our calculation. For example, generating δ^13^C_kerogen_ <-40‰ at 25°C necessitates *^13^ε_rub_* values >30‰. This is larger than any modern known rubisco (∼10-30‰; Supplemental Dataset 1). Although possible, current studies of reconstructed ancestral rubiscos find that, if anything, they actually exhibit smaller *^13^ε_rub_* values compared to extant rubiscos (17.2 vs. 25.2‰; (70)). Finally, a cyanobacterium expressing a similar ancestral rubisco (96.4% protein sequence identity to the ancestral rubisco in (70)) found *^13^ε_p_* values to be similar to that of its modern counterpart (*^13^ε_p_* values of ∼22-26‰ at 2% CO_2_; (63)). Therefore, current experiments are not consistent with ancestral *^13^ε_rub_* being larger than modern values and are thus challenging to reconcile with (very) hot Archean oceans under the assumption that rubisco was the dominant carbon fixing enzyme.

Overall, our analysis of the organic carbon record leads to the following proposals: (*i*) Over the Phanerozoic, isotope effects of photoautotrophs were either smaller than typically assumed or controlled in part or whole by CO_2_ limitation or CCMs; (*ii*) For the Proterozoic, we find that temperatures were either somewhat warmer than modern (40-50°C), *^13^ε_rub_* again smaller than what is typically assumed, or that *^13^ε_p_* was set by CCMs and/or CO_2_ limitation. We note that the latter may be somewhat surprising given the typical assumption that, outside of global glaciations, *p*CO_2_ was significantly higher than today (e.g., (71)); (*iii*) For the Archean, the data is generally inconsistent with some reconstructions of very high temperatures (up to 80°C).

### 3.4 Rubisco Biochemistry

A long-standing question of rubisco biochemistry is why at 25°C, *^13^ε_rub_* varies by over 20‰ at across various organisms. A leading hypothesis is that this variation is controlled by the same enzymatic properties that also cause variations in specificity (*S_C/O_*; (14)), which is the ratio of the catalytic efficiency of carboxylation (*V_C_/K_C_*) vs. oxygenation (*V_O_/K_O_*) (72). *V* and *K* are the Michaelis constants for RuBP-saturated reaction with CO_2_ and O_2_ respectively. This hypothesis was originally supported by the observation that *^13^ε_rub_* linearly correlates with *S_C/O_* (5 species, *P*=0.04, *R^2^* = 0.80) while the oxygen kinetic isotope effect (^18^KIE) does not (3 species, *P*=0.5; *R^2^* = 0.42). This analysis included both Form I and II rubiscos from plants and bacteria. (14) proposed that this relationship results from degree to which the transition state for CO_2_ addition to RuBP resembles the product vs. the reactant. Specifically, the more product-like the transition state (lower transition-state-energy), the more tightly the CO_2_ is bound and thus the higher its specificity and larger the *^13^ε_rub_*.

Since the initial study of (14), additional *^13^ε_rub_* values have been measured across diverse organisms at ∼25°C and the correlation is no longer apparent (14 species; *P*=0.34, *R^2^* = 0.075; Fig. 6A), challenging the initial proposal. Additionally, the model of (14) assumed that the carboxylation of RuBP is irreversible, but if this assumption is relaxed, one can arrive at different biochemical interpretations of *^13^ε_rub_* (e.g., see discussions in (15–18)). Based on this, we consider it unresolved from a biochemical perspective why *^13^ε_rub_* varies among rubiscos.

**Figure 6:**
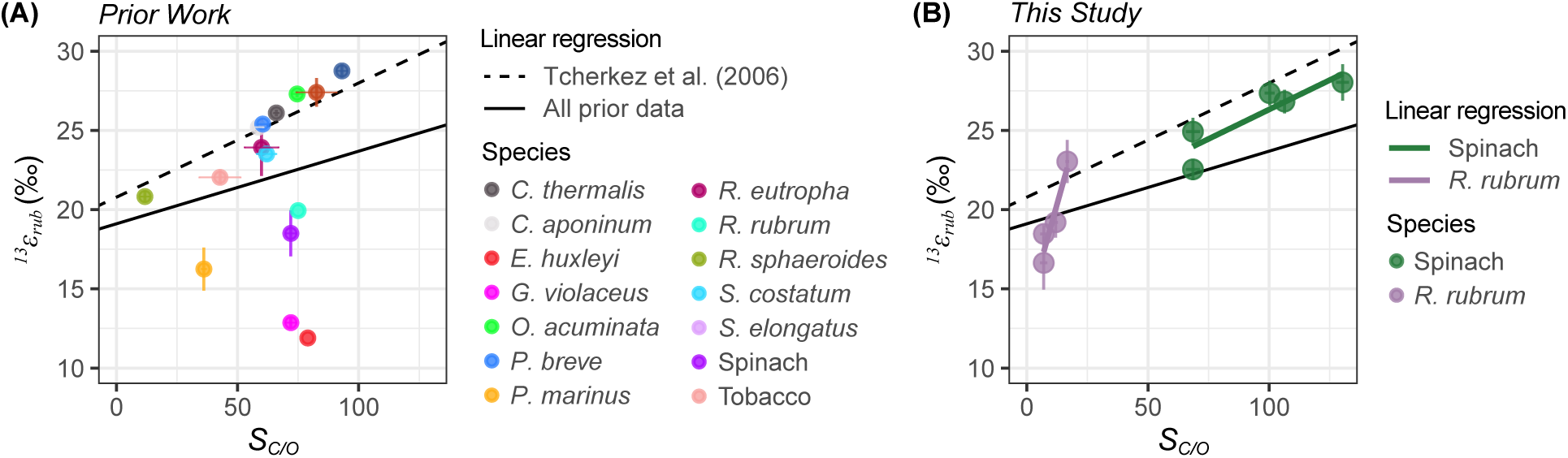
Rubisco carbon kinetic isotope effect and specificity data. **(A)** Prior *^13^ε_rub_* values of wild-type rubiscos from *in vitro* experiments performed at ∼25°C. See Supplemental Dataset 1 for compiled literature values, references, and more details. When the same species was measured across multiple studies, individual measurements across all studies were averaged. When possible, paired *^13^ε_rub_* and *S_C/O_* are taken from the same study, otherwise *S_C/O_* data was sourced from the compilation of (86). Linear regression from (14) is shown as a dashed line; solid line shows linear regression through all prior data. Uncertainties on data points are as reported from the original study or represent the standard deviation of *^13^ε_rub_* averaged across multiple studies. **(B)** Spinach and *R. rubrum* data from this study. Temperature-dependent changes in *S_C/O_* are taken from prior studies (87–89); Figure S4). Uncertainties on data points are ±1SE or smaller than the plotted circle. Uncertainties on all linear regressions are not shown for figure clarity.

Prior work has generally looked for the biochemical mechanism of *^13^ε_rub_* by empirically comparing it to *S_C/O_* or other enzymatic parameters measured at 25°C. 25°C is presumably used for this comparison because it is the typical temperature at which all the various enzymatic and isotopic parameters are measured. However, it is well known that rubisco carboxylation, oxygenation, and thus *S_C/O_* parameters are all strongly temperature dependent (see (73)). This matters as both the absolute and relative change in *S_C/O_* for the same temperature change are not constant between different rubiscos. For example, for the rubiscos examined here, spinach’s *S_C/O_* decreases in absolute value by 64 (133 to 69) from 10-35°C, compared to an absolute decrease of 10 (17 to 7) for *R. rubrum* (Fig. S4). For spinach and *R. rubrum*, both *S_C/O_* and *^13^ε_rub_* decrease as temperature increases, creating a positive correlation of *S_C/O_* and *^13^ε_rub_* (Fig. 6B). However, the slopes are significantly different as the absolute decrease in *S_C/O_* is smaller for *R. rubrum* even though its decrease in *^13^ε_rub_* is similar to spinach (∼4.5‰; Fig. 6B).

This leads to the question of how much observed correlations in the KIEs and enzyme kinetics of rubisco have been created or obscured by using 25°C as the comparison point. For example, we consider it an open question as to whether various relationships be normalized to 25°C or to the physiologically relevant temperature that an organism grows at in its natural environment. The optimum temperature of rubisco’s maximum carboxylation rate (*V_C_*) varies by species, and this temperature is often correlated with the organism’s growing environment (22). An alternative approach would be to examine the underlying thermodynamic variables that set the kinetic and isotopic parameters. For example, the temperature dependence of *S_C/O_* is largely set by the difference in enthalpy of free CO_2_ vs. O_2_ vs. in the transition state (ΔH^‡^), whereas absolute differences at a given temperature are related to differences in entropy of the reactants vs. the transition state (ΔS^‡^; (14, 74, 75)). In comparison, the temperature dependence of isotope effects originates, in the simplest of kinetic explanations, from the difference in activation energies of substrate vs. transition states (76, 77). Follow-up work that measures the temperature-dependence of *^13^ε_rub_* from additional species may help develop a stronger mechanistic understanding of the factors that drive rubisco’s kinetic isotope effects.

## 4. Conclusion

We have shown that the *^13^ε_rub_* of spinach and *R. rubrum* rubiscos are temperature dependent with statistically indistinguishable slopes, despite being from phylogenetically distinct enzyme and species clades. The observed temperature dependencies are large (∼4.5‰ over 25°C) and are of similar size to variations in δ^13^C_org_ and *^13^ε_p_* observed in modern terrestrial and marine environments. Therefore, the temperature-dependence of rubisco’s ^13^KIE is of sufficient magnitude to be considered as a variable that affects the δ^13^C of plants, algae, and organic carbon on modern and ancient Earth. Finally, from a biochemical perspective, our results raise an important question as to what temperature the isotopic and non-isotopic parameters of rubisco should be compared.

## 5. Materials and methods

### 5.1 Rubisco purification and acquisition

Rubisco from *R. rubrum* was expressed heterologously and purified from *E. coli* following established methods (78–80). Briefly, the *R. rubrum cbbM* gene (UniProt: P04718) was cloned into a pET28 vector (79) with an N-terminal His_14_-bdSUMO engineered protein tag (81) and transformed into chemically competent BL21 DE3 Star competent *E. coli* cells (MacroLab, Berkeley, USA). Expression of the pET28 vector was induced with IPTG (isopropyl β-D-1-thiogalactopyranoside), cells were lysed at via high pressure homogenization (Emulsiflex-C3, AVESTIN, Inc., Ottawa, Canada), and then rubisco was purified via poly-histidine affinity tag attachment to a nickel bead column (HisPur Ni-NTA resin, Thermo Fischer) and subsequent cleavage with SUMOlase enzyme. SUMOlase enzyme was prepared as previously described in (78). Rubisco purity was checked via SDS-PAGE (Fig. S5) and enzyme was stored with 25% glycerol in our reaction buffer (described below) at -80°C until used in the KIE assay measurement. See Supplemental for more detail. Spinach rubisco was obtained as lyophilized powder (Sigma-Aldrich R8000; CAS Number 9027-23-0) and contains other components seen in fresh, spinach lysate (Fig. S5). See Supplemental for more detail.

### 5.2 KIE Assays

All reactions were performed in “KIE buffer” at pH 8.0 (50 mM Bicine, 25 mM MgCl_2_, 10 mM HCO_3_^-^, 1 mM DTT). Experiments were initiated by activating rubisco in KIE buffer for ∼20 min at the reaction temperature (10, 22 or 35°C). Following activation, carbonic anhydrase (CA) was added (bovine erythrocyte; Sigma, C3934-500MG) followed by high-purity RuBP (≥99.0%; Sigma, 83895) dissolved in 50 mM bicine. For the spinach KIE assays, the amount of rubisco and CA varied from 5-68 mg and 0.2-3 mg respectively. For the *R. rubrum* assays 0.2-1 mg of rubisco and 0.4-3 mg of CA was used. See Table S2 for exact reaction conditions of each assay.

Next, the solution was added into a 10 mL gas-tight syringe (Hamilton 1010W, 81610) fitted with a luer septum adapter (Hamilton 31335) and a high temperature, low-bleed septum (Hamilton 75810). The inside of the syringe was manually beveled (by us) to form a divot to gather gas bubbles which were then ejected. After the solution was added to the syringe, the reaction was considered isolated from the atmosphere and to have begun. The time between addition of RuBP and the first sampling immediately following isolation in the syringe was ∼2-3 minutes. This first sampling is our initial time point. Over these 2-3 minutes, some reaction occurs such that the concentration and isotopic composition of our time zero is modified from that of the initial KIE buffer.

Next, sample aliquots of the KIE buffer were taken and quenched as follows. We subsampled 100-200 μL aliquots from the 10 mL syringe using a 250 μL gas-tight syringe for fluids (Hamilton 1725) fitted with a 3.5-inch, 22s gauge needle (Hamilton 7806-02). We always took and discarded an initial aliquot to clear the sampling syringe prior to collection for measurement of isotopic compositions and DIC concentration, and the sampling syringe was rinsed three times with deionized water between samples. In a given experiment, the same volume is sampled every time (i.e. a 200 μL sample is taken every time) so that changes in the measured amount of total carbon only reflected changes in concentration rather than changes in total volume. The solution was then immediately injected into a prepared round-bottom 12 mL borosilicate glass vials (Labco Exetainer E2863) sealed with either single or double-wadded caps; the cap was the same within each experiment. These tubes were prepared in advance with 200-300 μL of phosphoric acid (85% w/v) that were degassed for 25-30 seconds with He gas (Grade 5.0); the amount of phosphoric acid used was always the same for a given experiment.

Reaction temperatures were controlled as follows. Reactions at 10°C were conducted inside an oven with thermostat control placed inside a cold room (∼4°C). Reactions at 35°C were conducted in a thermostat-controlled room. Reactions at 22°C were conducted on the benchtop. For all experiments, all materials (KIE Buffer, Exetainer tubes, sampling syringes) were allowed to thermally equilibrate overnight at the experimental temperature prior to use, and temperatures were monitored independently with the same mercury thermometer across experiments.

### 5.3 Carbon Isotope Measurements

CO_2_ isotopic compositions were determined with a continuous-flow isotope ratio mass spectrometer (CF-IRMS; ThermoFisher Scientific Delta V Plus interfaced with a ThermoFisher Scientific GasBench II) at the UC Berkeley Center for Stable Isotope Biogeochemistry. Isotopic compositions are reported using delta notation (δ^13^C). All values are reported on the VPBD scale based on an assigned composition of the working reference gas. Each analytical run was composed of a series standard and sample measurements as follows: 1 gas standard, 1 blank (He 5.0 gas in sample bottle with phosphoric acid), four concentration calibration samples, and then six samples. Repeated measurements of various standards were performed to evaluate if any drift occurred over the course of a run. In no cases was measurable drift observed. For all measurement sessions, we tested for instrument linearity (measurable dependence of sample amount vs. isotopic composition) by measuring varying volumes of our KIE Buffer treated identically to samples as above. In all cases, we did not observe a measurable dependence. In some runs, two carbonate standards with a 38‰ difference were measured to independently check instrument linearity across this range. In all cases, the mean and standard deviation were within 1‰ of the expected difference. Samples with measurable air contamination were discarded.

### 5.4 Relative total inorganic carbon determinations

Changes in DIC concentrations were determined using the measured mass 44 peak area (corresponding to ^12^CO_2_) as a proxy for the total carbon in a measurement. For a given analytical run in which an isotope effect is being determined, we injected the same volume of liquid into identical Exetainer tubes pressurized in the same manner with the same amount of phosphoric acid. As such, any measured differences in the mass 44 peak area correlate to differences in the total number of moles of CO_2_ in the headspace and thus in the original DIC. For each measurement, we constructed calibration curves correlating the instrumental response of mass 44 peak area vs. DIC (Fig. S6). This was done by taking the initial KIE buffer (with no enzymes or RuBP added) and injecting known volumes into Exetainers with phosphoric acid prepared identically as the samples. In doing this, we measure mass 44 peak areas across a range that spans the total DIC measured in sample as a function of time. We then performed a linear regression of measured mass 44 peak area vs. volume of KIE Buffer and used this to normalize measurements of mass 44 peak area vs. the initial sample time point to changes in DIC. Reponses over this range were determined in all cases to be linear. Note that we do not attempt to quantify actual DIC concentrations directly as such are not needed for measurements of isotope effects using the Rayleigh distillation approach.

### 5.5 Determination of isotope effects

We determined values of *^13^ε_net_* by fitting data to a linear line for *^13^R_t_/^13^R_i_* (y-axis) vs ln(*f*) (fraction DIC remaining vs. initial; x-axis) Figs. S7-S8). *^13^R* is calculated as (1000 + δ^13^C_t_)/(1000 + δ^13^C_i_), where δ^13^C is the average δ^13^C of the CO_2_ of the acidified DIC pool at each timepoint (*t*) vs. the initial timepoint (*i*). For our experiments, we aim to have total DIC consumption ≥50% relative (Table S2) in order to generate robust variations in *^13^ε_net_* vs. *f*. We did this because the parameter with the most measurement uncertainty is *f*, especially when changes in concentration are small. A variety of fitting procedures have been proposed for the determination of *^13^α_rub_* including point by point determinations (e.g., (39)). Here, we used all data together to fit the slope.

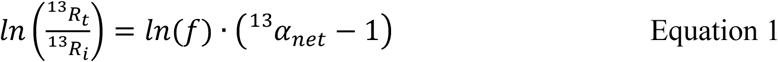

Values of *^13^*α were solved for and then *^13^*ε was calculated. For a system following the Rayleigh distillation model, the slope of line in Equation 1 equates to (*^13^α_net_* - 1), which is equivalent to - *^13^ε_net_*. *^13^α_net_* incorporates all isotope effects occurring during the removal of CO_2_ from the system including those from rubisco and DIC equilibrium. At the pH of our experiments (∼8), 98% of the DIC is HCO_3_^-^ such that we can approximate that the DIC measured effectively represents the isotopic composition of HCO_3_^-^ in solution, as done by prior studies (e.g., (43)). We therefore defined the equilibrium isotope effect as *^13^α_CO2(aq)-HCO3-_*. This equilibrium isotope effect corresponds to *^13^ε_equil_* in epsilon notation. Carbonic anhydrase rapidly catalyzes this equilibration such that equilibrium of carbon isotopes between the DIC species is maintained at all times (82).

We related the various fractionation factors as follows:

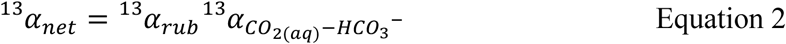

To calculate *^13^α_rub_*, we took the measured value of *^13^α_net_* and corrected it for the value of *^13^α_CO2(aq)-HCO3-_*, which is itself temperature dependent. We calculated *^13^α_CO2(aq)-HCO3-_* by combining experimental calibrations for the fractionations for CO_2(aq)_ and CO_2(g)_ and for HCO_3_^-^and CO_2(g)_ from (40):

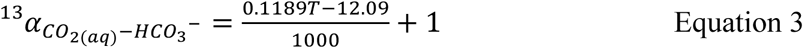

Here, *T* is in Celsius. We also note that some prior work (e.g., (38, 83)) applied a correction based on pH. Applying this correction to our data increased all KIE values by ∼0.2‰ and, as such, did not affect the interpretation of our results.

### 5.6 Estimates of terrestrial and marine temperatures

For terrestrial environments, we used estimated mean growing season temperatures (*T_gs_*) of plants as a proxy for average photosynthesis temperatures on land, taken from the global leaf data set of (24). We filtered the data to remove samples >2,000 m in order to remove elevation effects on latitudinal temperature gradients. Finally, a latitudinal gradient was fit using a LOESS curve. For marine systems, we calculated a weighted sea-surface temperature (SST_w_) where SSTs from 2013-2022 from the NOAA ERSST dataset (54–56) were weighted by net primary productivity (NPP) calculated using the Carbon-based Production Model from (51) (Fig. S2).

SST_w_ for each year was calculated as a weighted mean where, at each latitude, the monthly SST was weighted by the corresponding NPP value. A final weighted SST_w_ mean was then calculated at each latitude, across all longitudes, across all ten years.

## 6. Acknowledgements

We acknowledge Wenbo Yang for kind assistance with measuring samples and Phoebe Lam for help with accessing GEOTRACES data. Work was funded by NASA Exobiology Grant NNH20ZDA001N-EXO to D.A.S and P.M.S. and a Packard Fellowship from the David Lucile Packard Foundation to P.M.S. and A.K.L. R.Z.W. is an Agouron-Resnick Fellow of the Life Science Research Foundation.

