## Supplementary text for "Form I and II Rubiscos Exhibit Temperature Dependent Carbon Kinetic Isotope Effects"

**This PDF file includes:**

Supporting text
Figures S1 to S8
Tables S1 to S3
Legends for Datasets S1 to S2
SI References

**Other supporting materials for this manuscript include the following:**

Datasets S1 to S2

**Supporting Information Text**

**Table of Contents**

1) *R. rubrum* rubisco transformation and expression in *E. coli*
2) *R. rubrum* rubisco purification
3) Figures
4) Tables
5) Dataset S1 and S2 descriptions
6) References

**Figure List**

Figure S1: Results plotted as $ln(^{13}\alpha_{rub})$ vs. 1/T
Figure S2: Example workflow calculating NPP-weighted SSTs for 2019.
Figure S3: Comparison of variations in the carbon isotope composition of particulate organic carbon vs. predicted carbon KIE with latitude.
Figure S4: Temperature-dependence of Specificity for *R. rubrum* and spinach.
Figure S5: SDS-PAGE Gel of *R. rubrum* and Spinach rubisco enzymes.
Figure S6: Calibration curves.
Figure S7: Rayleigh curves for *R. rubrum* experiments
Figure S8: Rayleigh curves for Spinach experiments

**Table List**

Table S1: Results
Table S2: Reaction conditions
Table S3: Linear regressions for $\ln(^{13}\alpha)$ vs. 1/T.

**1) *R. rubrum* rubisco transformation and expression in *E. coli***

The *R. rubrum cbbM* gene (UniProt: P04718) was cloned into a pET28 vector (*1*) with an N-terminal $His_{14}$-bdSUMO engineered protein tag (*2*) and kanamycin resistance that was synthesized by Twist Biosciences. The pET28 plasmid was then transformed into BL21 DE3 Star competent *E. coli* cells (MacroLab, Berkeley, USA) via heat shock, revived in LB (Luria-Bertani) at 37°C while shaking at 200 RPM (revolutions per minute) for ~1.5 hours, and then plated for single colonies on LB agar. Single colonies were then grown overnight to high densities in 10 mL of LB with kanamycin at 1,000x dilution at 37°C and 200 RPM. This 10 mL growth was then used to inoculate a 1 L flask of LB with kanamycin at 1,000x dilution; cells were then grown to mid-log phase at 37°C to an optical density at 600 nm ($OD_{600}$) of ~0.6. At $OD_{600}$ ~ 0.6, expression of the pET28 vector was induced with 1 mM IPTG (isopropyl β-D-1-thiogalactopyranoside) and then flasks were incubated overnight at 16°C at 180 RPM. Next, cells were pelleted at 5,000xG at 4°C for 20 minutes then resuspended in 20 mL of lysis buffer (50 mM sodium phosphate, 300 mM NaCl, 10 mM imidazole, 5% glycerol, 2mM $MgCl_2$) with 1 mM PMSF (phenylmethylsufonyl fluoride) to inhibit native protease activity. Cells were then frozen at -80°C until cell lysis.

**2) *R. rubrum* rubisco purification**

All work was performed on ice or at 4°C. Cells were thawed and lysed via high-pressure homogenization with an Emulsiflex-C3 (AVESTIN, Inc., Ottawa, Canada) and cell lysate was incubated with approximately 1 mg/mL of DNase I (REF 10104159001; Roche) at 4°C for ~1 hour. This cell lysate was centrifuged at 15,000 x *g* for 20 minutes at 4°C and the pellet was removed. The soluble fraction was filtered using 0.44 µm filters and incubated with HisPur Ni-NTA resin (Thermo Fischer) with gentle rocking for 1 hour at 4°C to batch bind the $His_{14}$-bdSUMO tag to the resin. Prior to batch binding, the resin was first washed with an equilibration buffer ("Ni Equilibration Buffer"; 50 mM sodium phosphate, 300 mM NaCl, 10 mM imidazole, 10% glycerol). After batch binding, the total solution was placed into a column and washed twice, first with a 25 mM imidazole wash buffer (50 mM sodium phosphate, 300 mM NaCl, 25 mM imidazole, 10% glycerol), and then a 50 mM imidazole wash buffer (50 mM sodium phosphate, 300 mM NaCl, 50 mM imidazole, 10% glycerol). Next, washed beads were incubated with the SUMOlase enzyme in SUMOlase buffer at pH 8.0 (20 mM HEPES, 100 mM NaCl, 15 mM imidazole, 20 mM $MgCl_2$) and rocked gently overnight at 4°C. SUMOlase enzyme was prepared as previously described in (*3*); briefly, the SUMO-specific protease from *Brachypodium distachyon* (bdSENP1) with a 7x His-Tag was expressed by transforming the pSF1389 plasmid into chemically competent BL21 DE3 Star *E. coli cells* (Macrolab), lysed via high pressure homogenization, bound to Ni-NTA Resin (Thermo Fischer), and cleaved with the TEV protease (MilliporeSigma).

After overnight incubation with the SUMOlase enzyme, the column was washed an additional time with SUMOlase buffer and the total flow through was centrifuged at 14,000 x *g* for 7 minutes at 4°C. Rubisco protein concentration was determined via absorption at 280 nm with the NanoDrop One$^{C}$ (Thermo Fischer) and purity was determined via SDS-PAGE. Next, crude protein was concentrated via centrifugation with centrifugal filters (30 kDa Amicon Ultra filters, Merck Millipore) and stored at 25% glycerol at -80°C until the KIE Assay measurement. In general, we obtained ~4 mg of crude protein from 1 L of growth.

**3) Figures**

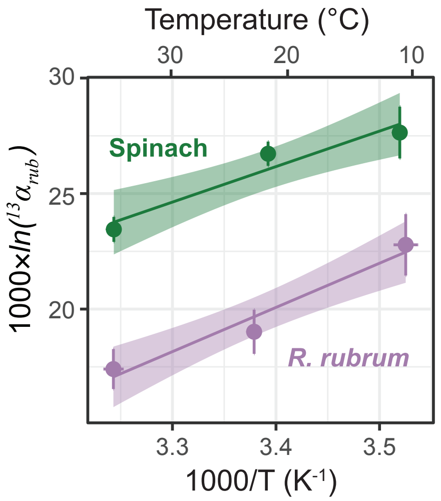


**Figure S1: Results plotted as $\boldsymbol{ln(^{13}\alpha_{rub})}$ vs. 1/T.** 1000×$ln(^{13}\alpha_{rub})$ vs. 1000/T for spinach (green) and *R. rubrum* (purple); replicates at 20-22°C and 35°C were averaged before fitting. Fits show linear regressions with 68% c.i.; see Table S3 for linear regression results.

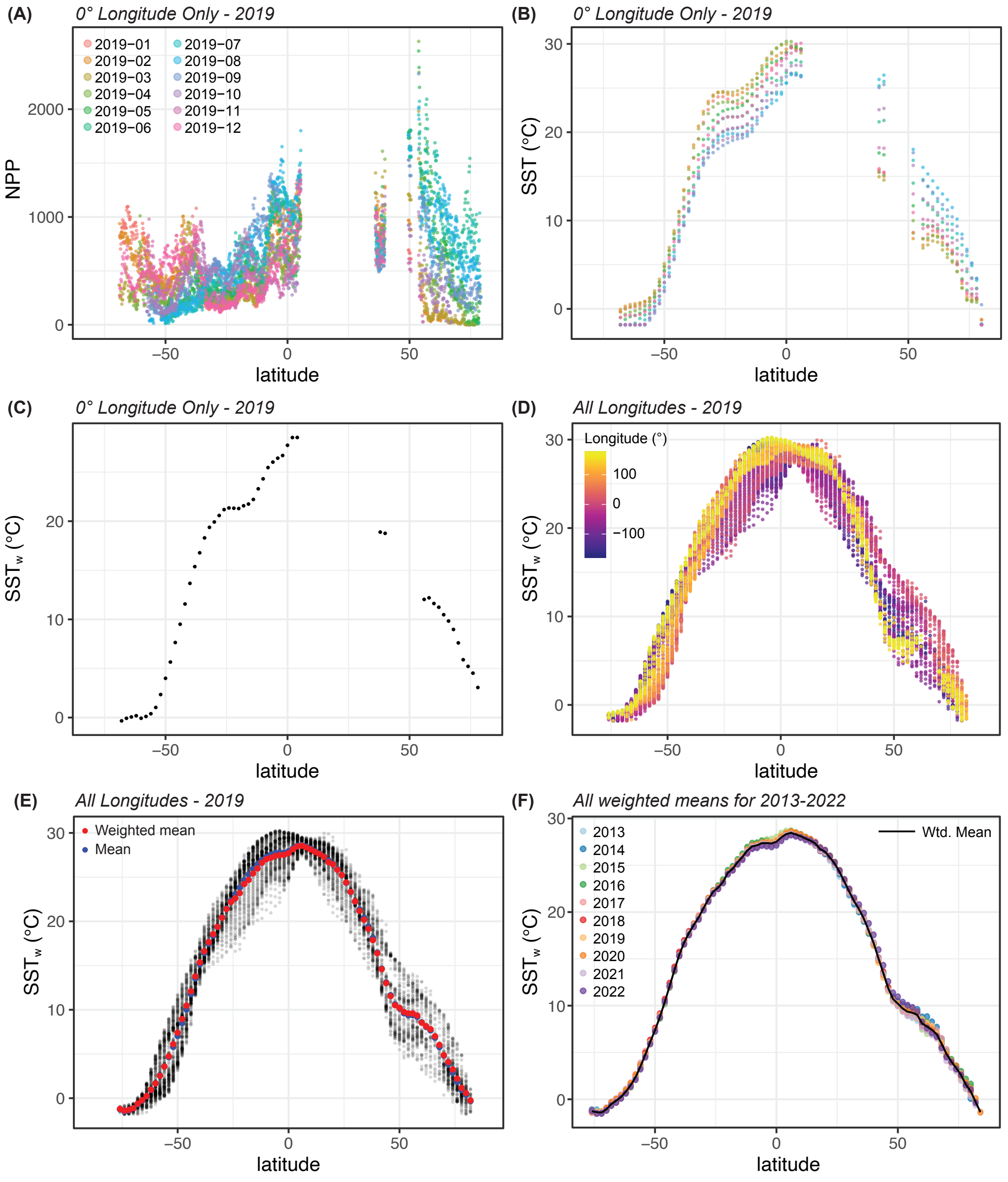


**Figure S2: Workflow calculating NPP-weighted SSTs using 2019 as an example. (A)** Net primary productivity (NPP) data from (*4*) (based on the Carbon-based Production Model; CbPM). Data is for the year 2019 at a longitude of 0° only. Colors indicate year and month (yyyy-mm). Data is gridded at a 0.167° resolution. **(B)** Sea surface temperature (SST), at a 2° resolution, from the NOAA ERSSTv5 dataset (*5*) from 2019, and also at a longitude = 0°. Colors are the same as in panel A. **(C)** NPP-weighted SST ($SST_w$) calculated for 2019 at longitude = 0°. $SST_w$ was calculated as a weighted mean in which the monthly SST was weighted by the corresponding NPP value. NPP data was resampled to the coarser SST grid prior to weighting using bilinear interpolation via the R package *terra* (v.1.8.93; *resample()* function (*6*). **(D)** $SST_w$ for 2019 now shown for all longitudes, indicated by color. **(E)** Black points are the same data as in panel (D). Red circles show a weighted $SST_w$ mean calculated at each latitude, across all longitudes, while blue circles show a simple mean. **(F)** Weighted means, calculated in the same manner, for 2013-2022 shown in colors. The black line, shown in the main text, shows the weighted mean of all years.

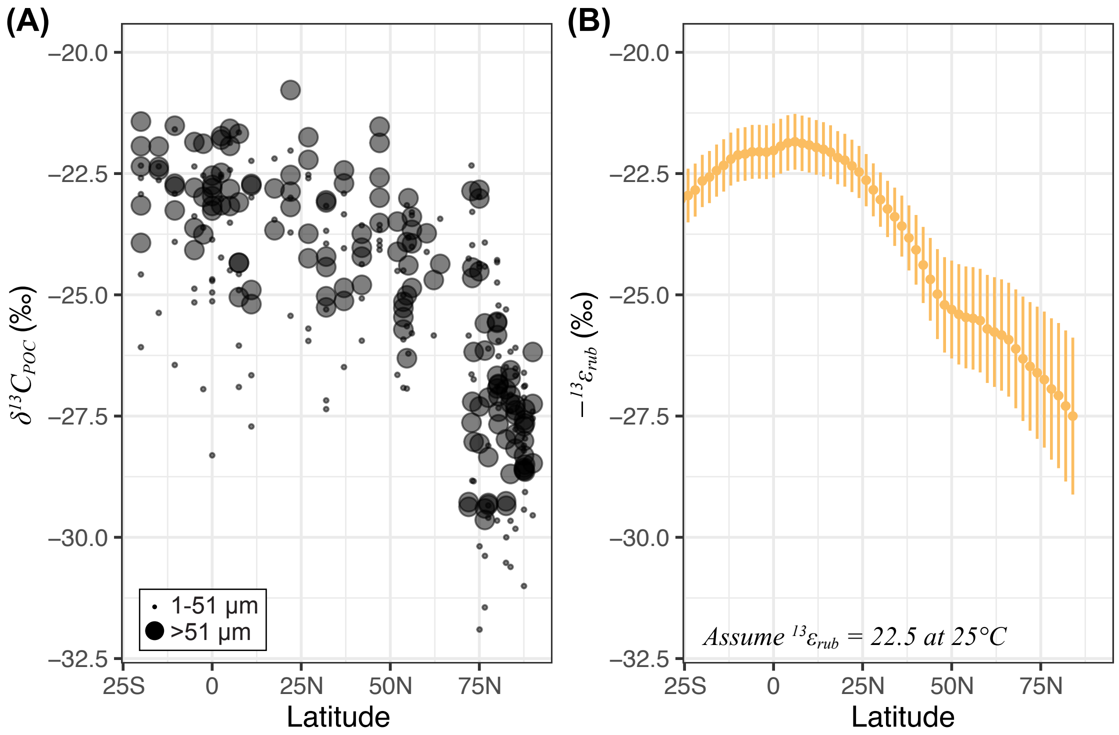


**Figure S3: Carbon isotope composition of particulate organic carbon vs. predicted carbon KIE with latitude. (A)** Ocean POC carbon isotope data ($\delta^{13}C_{POC}$) was compiled from two GEOTRACES cruises – GN01 Arctic GEOTRACES cruise (Bering Sea to North Pole) and the GP15 Pacific Meridional transect cruise (Gulf of Alaska to Tahiti) (*7*–*9*). The GN01 cruise contained data from 60 to 72°N Latitude, while GP15 was from -20 to 54°N. Only data flagged with 1 (good), 2 (probably good), and 5 (changed; data adjusted during quality control) were plotted. Data were also filtered for a depth of less than 200 m as an approximation for the the photic zone. Small black circles indicate particles between 1-51 µm in size, while large black circles are for those larger than 51 µm. **(B)** Calculated variations in $^{13}\varepsilon_{rub}$ using the mean weighted SST from 2013-2022 (Figure 1) for temperature, and assuming an $^{13}\varepsilon_{rub}$ of 22.5‰ at 25°C so that the data in panels A and B would have a similar intercept for ease of visual comparison. Choosing a larger value for $^{13}\varepsilon_{rub}$ (i.e., 30‰ at 25°C) would shift the curve down in Panel B (towards more negative values of $-^{13}\varepsilon_{rub}$) but show the same trend. The y-axis shows negative $^{13}\varepsilon_{rub}$ values ($-^{13}\varepsilon_{rub}$) to be comparable to $\delta^{13}C_{POC}$ data – for both panels, a more negative number indicates greater relative depletion in $^{13}C$.

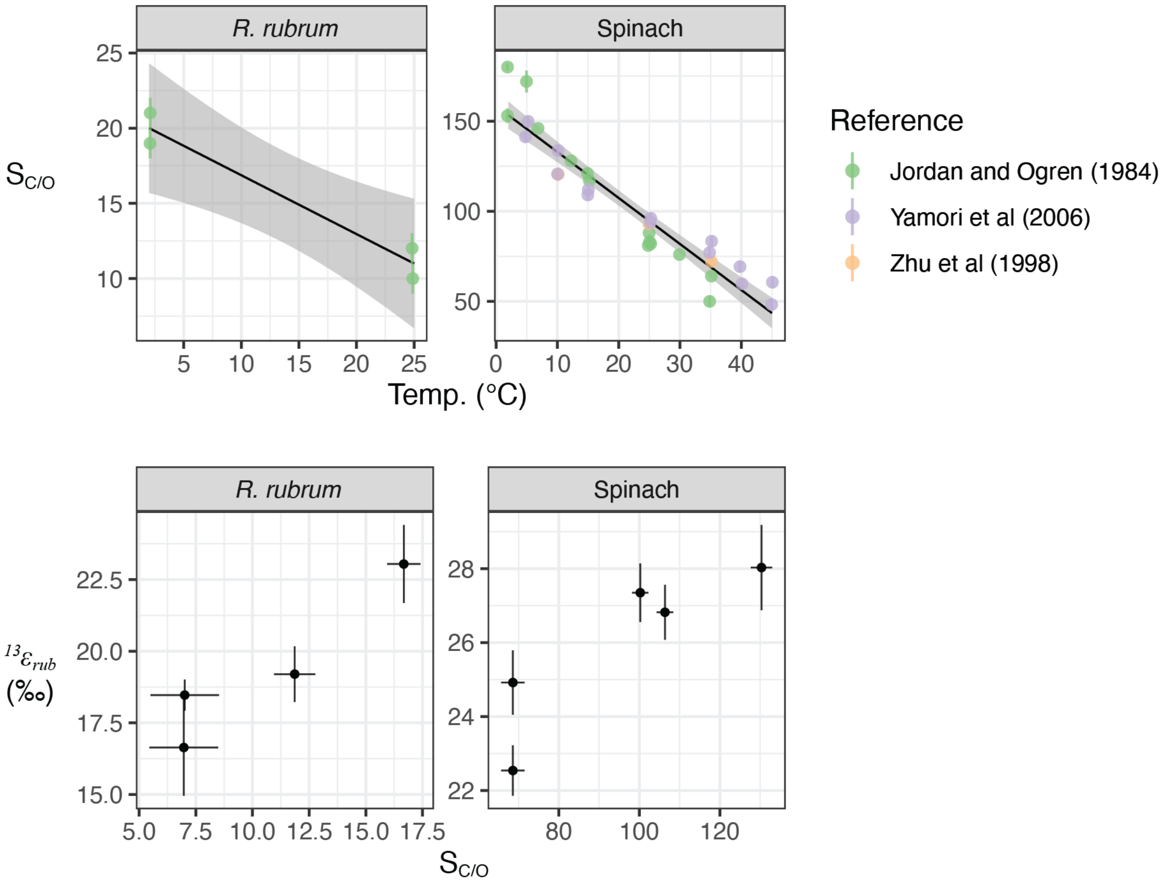


**Figure S4: Temperature-dependence of Specificity for *R. rubrum* and spinach.** Data is taken from (*10*) for *R. rubrum*, and from (*10–12*) for spinach. A linear regression was calculated through all points, and the corresponding $S_{C/O}$ value was calculated for our tested temperatures. Linear regression for *R. rubrum* was $S_{C/O}$ = (-0.39±0.06)T + (20.78±1.09) ($P$=0.02, $R^2$=0.953). Linear regression for spinach was $S_{C/O}$ = (-2.55±0.16)T + (158.43±3.95) ($P$=3.06×10$^{-16}$, $R^2$=0.903). Bottom panels show those predicted $S_{C/O}$ values (±1SE), predicted using the linear regression of $S_{C/O}$ vs. T from the top panels, paired with $^{13}\varepsilon_{rub}$ measured in this study; these data are plotted in Figure 6.

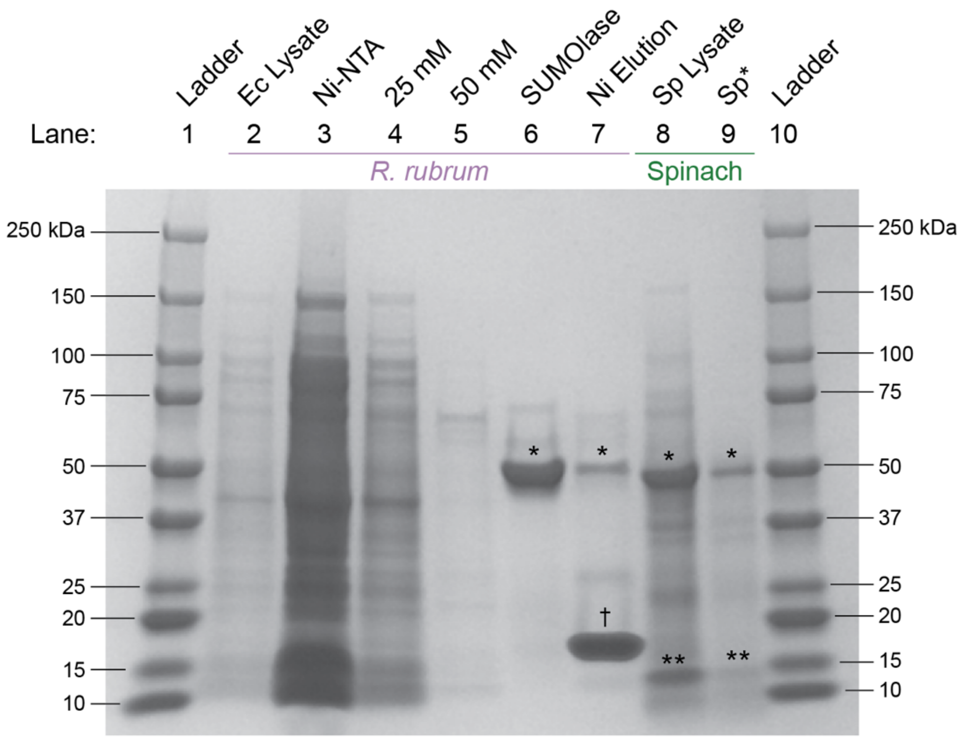


**Figure S5: SDS-PAGE Gel of *R. rubrum* and Spinach rubisco enzymes.** Gel shows typical purification results of *R. rubrum* from BL21 DE3 Star *E. coli* competent cells (Macrolab, Berkeley, CA) and of spinach rubisco obtained as lyophilized powder (Sigma-Aldrich R8000; CAS Number 9027-23-0). Variations in darkness across lanes reflect differences in protein elution concentrations. Concentrations (mg/mL) and purity (Absorbance at 260 vs. 280 nm; A260/A280) were determined via NanoDrop One$^{C}$ (Thermo Scientific) assuming 1 unit of absorbance at 280 nm = 1 mg/mL, using a baseline correction at 340 nm. Values are reported below in brackets [mg/mL; A260/A280]. Image above was adjusted for darkness, lightness, exposure, and contrast for viewing clarity. **Lanes 1 and 10:** Ladder from Precision Plus Protein Standards (Bio-Rad) showing molecular weights (kDa). **Lane 2:** "Ec Lysate" is filtered (0.45 µM), centrifuged whole cell lysate of *E. coli* [3.240 mg/mL; 1.97]. **Lane 3:** "Ni-NTA" is flowthrough (FT) after batch binding of *E. coli* lysate with HisPur Ni-NTA Resin (Thermo Scientific) [59.212 mg/mL; 2.03]. **Lane 4:** "25 mM" is FT after washing with 25 mM imidazole Ni wash buffer [9.561 mg/mL; 1.95]. **Lane 5:** "50 mM" is FT after washing with 50 mM imidazole Ni wash buffer [0.519 mg/mL; 1.88]. **Lane 6:** "SUMOlase" is FT after overnight incubation with 'SUMOlase' enzyme; star (*) is placed above the band for the *R. rubrum* rubisco large subunit monomer (RbcL) [0.978 mg/mL; 0.73]. This is the fraction used for KIE assays. **Lane 7:** "Ni Elution" is FT after washing with 300 mM imidazole Ni Elution buffer; dagger (†) is placed above the band for the polyhistidine 'SUMO' tag. Residual RbcL (*) is also seen. [0.131 mg/mL; 0.88]. **Lane 8:** "Sp Lysate" is whole cell lysate of spinach leaves bought fresh from a local grocery store. Leaves were frozen at -80°C, homogenized with mortar and pestle in a lysis buffer (20 mM HEPES, 50 mM NaCl, 5% glycerol, 2 mM $MgCl_2$, pH 8.0), rough filtered through a coffee filter, centrifuged (20 minutes, 15,000 xG, 4°C), and then filtered at 0.45 µM. Star (*) and double star (**) are placed above the bands for the large rubisco subunit (RbcL) and small rubisco subunit (RbcS) monomer bands respectively [7.195 mg/mL; 1.34]. **Lane 9:** "Sp*" is spinach rubisco obtained as lyophilized powder from Sigma-Aldrich re-suspended in deionized water [0.924 mg/mL; 1.71]. This was used for KIE assays.

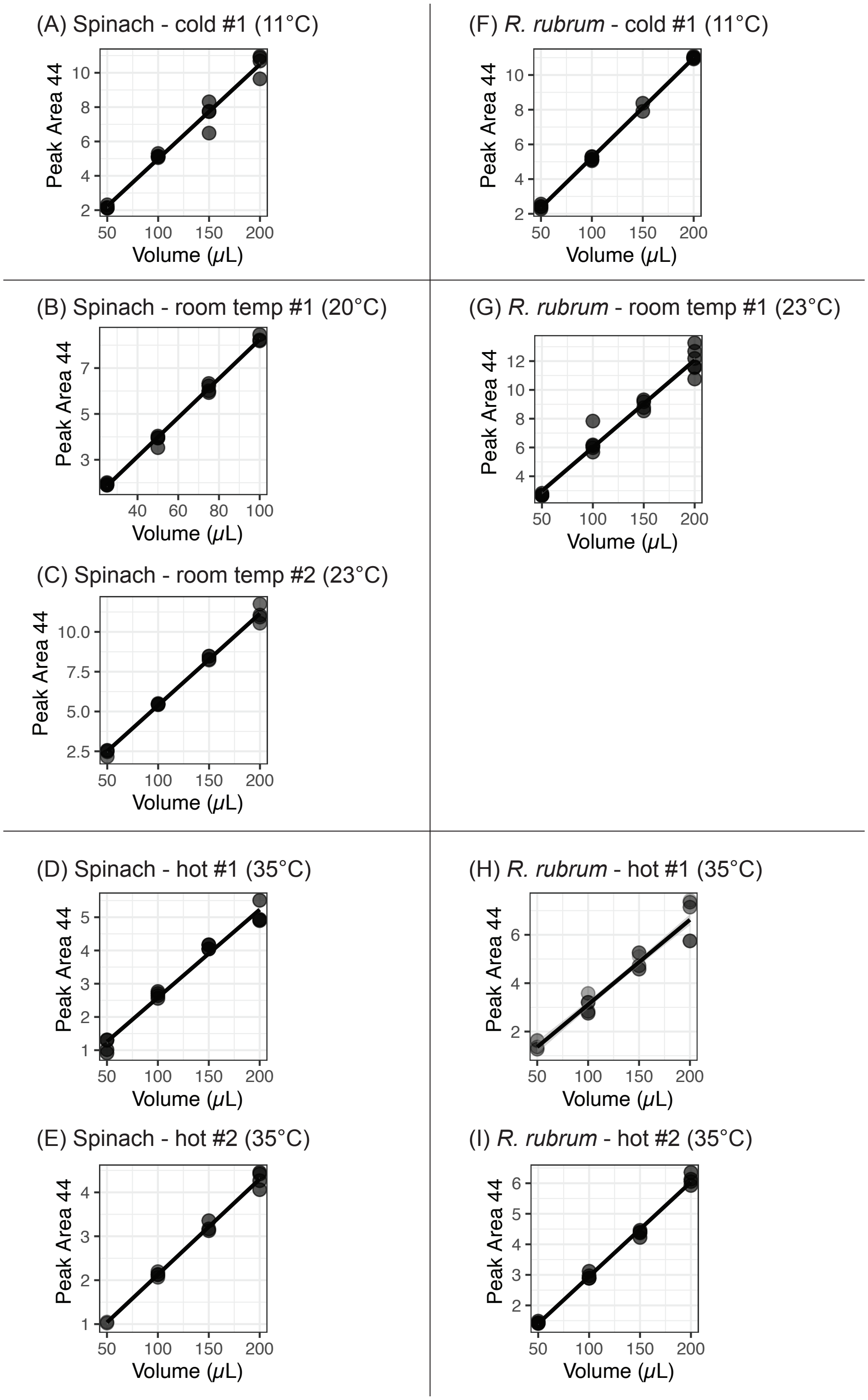


**Figure S6: Calibration curves.** *Spinacia oleracea* ("Spinach," A-E), *Rhodospirillum rubrum* ("Rubrum," F-I). Peak Area 44 indicates the first eluted mass 44 ($^{12}C^{16}O_2$) peak out of five total peaks; there is no measured uncertainty on this value. Volume (μL) is the volume of KIE buffer added. There are typically 4 replicates for each volume. 95% c.i. on linear regression is plotted above even though it is not readily visible.

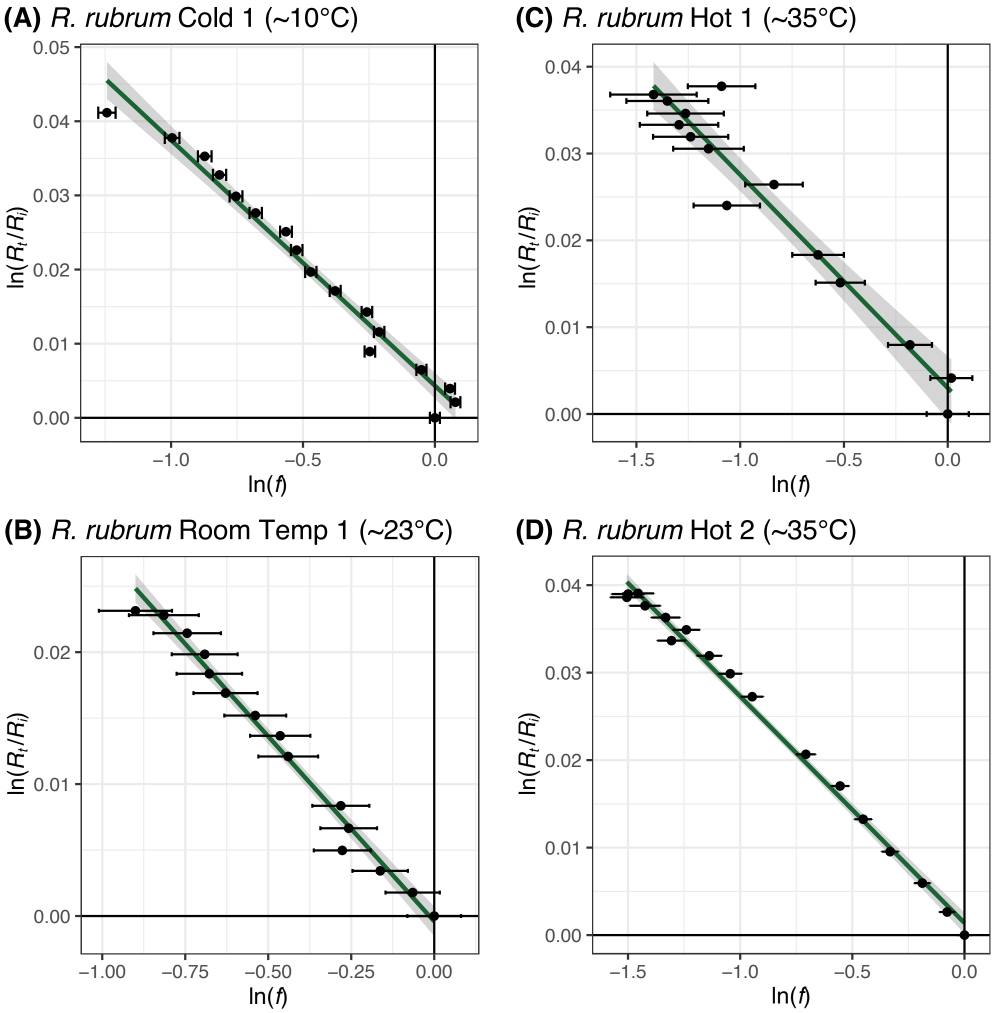


**Figure S7: Rayleigh curves for *R. rubrum* experiments.** Green line shows a linear regression while gray error envelope shows 95% c.i. Error bars on discrete data is ±1SE. Uncertainty on $ln(R_t/R_i)$ is plotted but is not visible as it is smaller than the marker. Error bars on discrete data were calculated from the calibration curve run with each experiment.

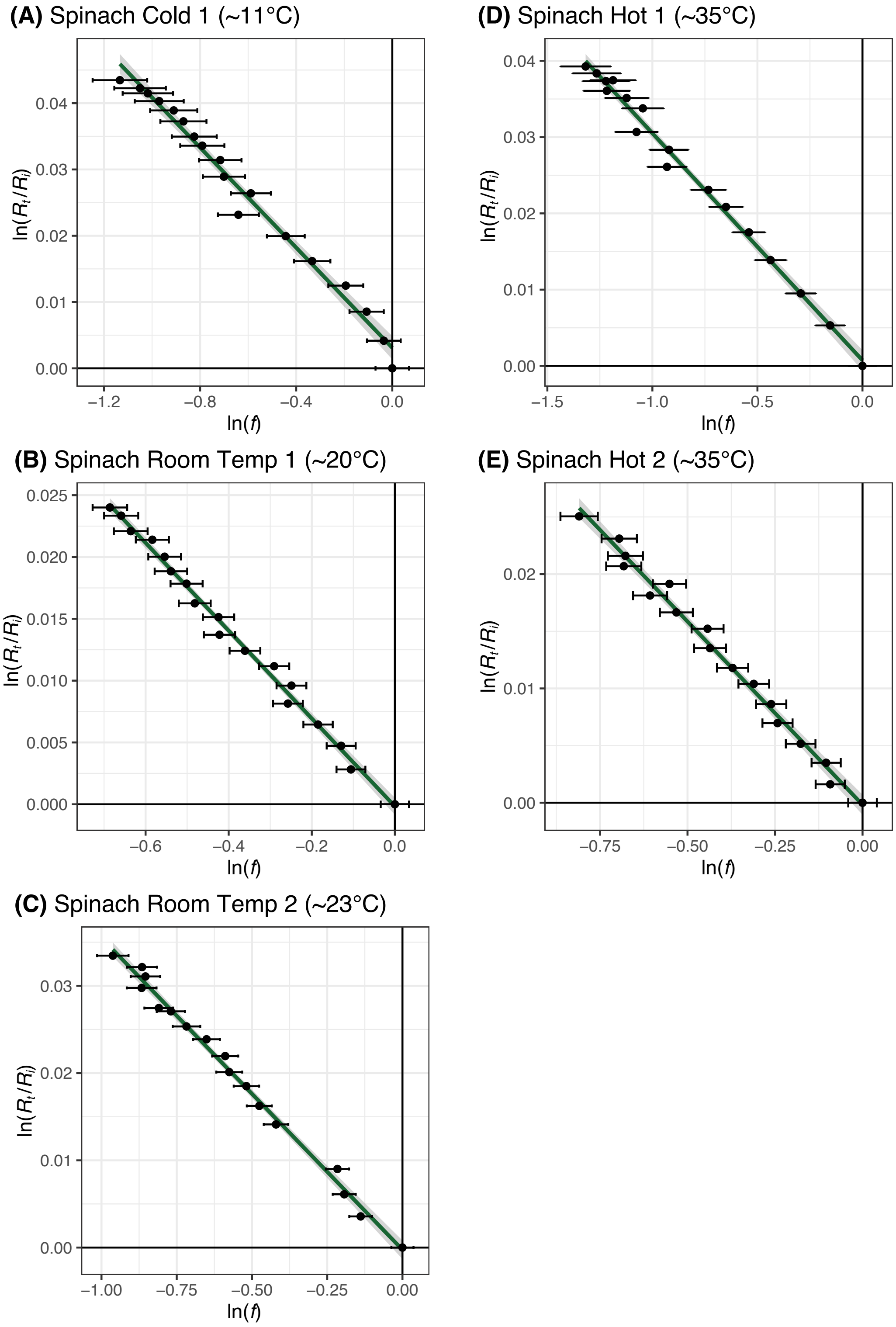


**Figure S8: Rayleigh curves for Spinach experiments.** Green line shows a simple linear regression while gray error envelope shows 95% c.i. Error bars on discrete data is ±1SE. Uncertainty on $ln(R_t/R_i)$ is plotted but is not visible as it is smaller than the marker. Error bars on discrete data were calculated from the calibration curve run with each experiment.

### 4) Tables

**Table S1: Results.**

| Strain | Temperature (°C) | | $^{13}\varepsilon_{net}$ | | | Adj. $R^2$ | $^{13}\varepsilon_{rub}$ | |
|---|---|---|---|---|---|---|---|---|
| | Mean | ±1SD | Estimate | ±1SE | *n* | | Estimate | ±1SE |
| Spinach | 11.0 | 0.4 | 39.23 | 1.15 | 18 | 0.984 | 28.03 | 1.15 |
| | 20.4 | 0.3 | 36.83 | 0.74 | 18 | 0.993 | 26.82 | 0.74 |
| | 22.8 | 0.2 | 37.08 | 0.79 | 17 | 0.992 | 27.35 | 0.79 |
| | 35.2 | 0.5 | 30.69 | 0.68 | 17 | 0.992 | 22.54 | 0.68 |
| | 35.2 | 0.5 | 33.09 | 0.87 | 17 | 0.988 | 24.92 | 0.87 |
| *R. rubrum* | 10.5 | 0.9 | 34.25 | 1.36 | 17 | 0.974 | 23.04 | 1.36 |
| | 22.8 | 0.4 | 28.85 | 0.97 | 15 | 0.983 | 19.20 | 0.97 |
| | 35.3 | 1.2 | 24.73 | 1.68 | 14 | 0.941 | 16.64 | 1.68 |
| | 35.2 | 1.4 | 26.59 | 0.54 | 17 | 0.993 | 18.47 | 0.54 |

These data are plotted in Figure 2 in the main text. Reported uncertainty on the net fractionation ($^{13}\varepsilon_{net}$) is derived from the linear regression of the Rayleigh curve, *n* indicates the number of data points fit and Adj. $R^2$ gives the goodness of fit. Fitting was performed in RStudio (v2023.06.1+524) with R (v4.4.0) [call: *lm()*] (*13*). The carbon KIE ($^{13}\varepsilon_{rub}$) is calculated after correcting for the equilibrium isotope effect (*14*); the uncertainty on this correction is negligible so the uncertainty on $^{13}\varepsilon_{rub}$ is the same as that for $^{13}\varepsilon_{net}$. SD = standard deviation. SE = standard error.

**Table S2: Reaction conditions.**

| Experiment name | Temp. (°C) | pH | Rubisco (mg) | Rubisco (mg/mL) | CA (mg) | CA (mg/mL) | Fraction consumed (*f*) |
|---|---|---|---|---|---|---|---|
| Spinach room temp 1 | 20.4 ± 0.3 | 8.00 | 5.0 | 1 | 0.24 | 0.05 | 50% ± 1% |
| Spinach room temp 2 | 22.8 ± 0.2 | 7.98 | 12.0 | 1.3 | 0.5 | 0.06 | 62% ± 1% |
| Spinach hot 1 | 35.2 ± 0.5 | 8.00 | 11.7 | 1.3 | 1.1 | 0.12 | 73% ± 1% |
| Spinach hot 2 | 35.2 ± 0.5 | 8.05 | 8.6 | 1.0 | 0.8 | 0.09 | 56% ± 1% |
| Spinach cold 1 | 11.0 ± 0.4 | 8.03 | 67.5 | 7.5 | 2.7 | 0.30 | 68% ± 2% |
| Rubrum room temp 1 | 22.8 ± 0.4 | 8.00 | 0.2 | 0.02 | 0.36 | 0.04 | 59% ± 2% |
| Rubrum hot 1 | 35.3 ± 1.2 | 8.00 | 0.4 | 0.04 | 1.4 | 0.16 | 67% ± 4% |
| Rubrum hot 2 | 35.2 ± 1.4 | 7.99 | 0.4 | 0.04 | 1.1 | 0.12 | 78% ± 1% |
| Rubrum cold 1 | 10.5 ± 0.9 | 8.00 | 1.0 | 0.1 | 2.5 | 0.28 | 71% ± 0% |

All reactions were performed in KIE buffer (50 mM Bicine, 25 mM $Mg^{2+}$, 10 mM $HCO_3^-$, 1 mM DTT, 10 mM RuBP) as described in the methods. pH was adjusted using HCl or NaOH. CA = Carbonic Anhydrase from bovine erythrocytes, used as lyophilized powder (Sigma, C3934-500MG).

**Table S3: Linear regressions for ln($^{13}\alpha$) vs. 1/T.**

| Strain | Fractionation | Slope ($m$) | | Intercept ($b$) | | Adj. $R^2$ |
|---|---|---|---|---|---|---|
| | | Estimate | ± 1SE | Estimate | ± 1SE | |
| Spinach | Net ($^{13}\alpha_{net}$) | 25.89 | 4.47 | -0.052 | 0.015 | 0.942 |
| | Rubisco ($^{13}\alpha_{rub}$) | 15.37 | 4.23 | -0.026 | 0.014 | 0.859 |
| *R. rubrum* | Net ($^{13}\alpha_{net}$) | 29.63 | 3.73 | -0.071 | 0.013 | 0.969 |
| | Rubisco ($^{13}\alpha_{rub}$) | 19.14 | 3.98 | -0.045 | 0.013 | 0.917 |

Best fits for $ln(^{13}\alpha) = m*(1/T) + b$, where temperature is in Kelvin. Reported uncertainty on $m$ and $b$ is ± 1SE from a simple linear regression. The covariance on the slope and intercept ($\sigma_{mb}$) is -0.633297. Fitting was performed in RStudio (v2023.06.1+524) with R (v4.4.0) [call: *lm*(); *vcov*()] (*13*).

**5) Dataset S1 and S2 Descriptions**

**Dataset S1 (separate file).** Literature compilation. Sheet 1 is a compilation of WT $^{13}\varepsilon_{rub}$ measured *in vitro* (i.e., from a pure enzyme assay) from (*15–29*) and Specificity ($S_{C/O}$) from (*3*, *15*, *24*, *30–34*). Values are labeled as follows:

| Column name | Description |
|---|---|
| Species or Strain | Species or strain of rubisco host. |
| Form | Rubisco clade. |
| Specificity: estimate | Reported Specificity ($S_{C/O}$) value; unitless. Values are taken as reported from each study, which typically reports the mean. Values with the superscript 'a' are the median value calculated from the compilation by (*33*). |
| Specificity: uncertainty | Reported uncertainty on Specificity ($S_{C/O}$); unitless. Values are taken as reported from each study, which typically reports the standard deviation. Otherwise, values in brackets are the 95% c.i. calculated from the compilation by (*33*). |
| Specificity: Reference | Reference from which Specificity is taken from. |
| $^{13}$KIE: estimate | Reported $^{13}\varepsilon_{rub}$ in units of per mil (‰). Values are taken as reported from each study, which typically reports the mean. Otherwise, values with the superscript 'a' are the reported median value. For *Rhodospirillum rubrum*, *Synechococcus elongatus*, and *Spinacia oleracea*, the estimate is the mean of all previously measured $^{13}$KIE values – see Sheet 2 for a compilation of those values. |
| $^{13}$KIE: uncertainty | Reported uncertainty on $^{13}\varepsilon_{rub}$ in units of per mil (‰). Values are taken as reported from each study, which typically reports the standard deviation. Otherwise, values in brackets are the 95% c.i. reported by that study. For *Rhodospirillum rubrum*, *Synechococcus elongatus*, and *Spinacia oleracea*, the uncertainty is the standard deviation of all previously measured $^{13}$KIE values – see Sheet 2 for a compilation of those values. |
| $^{13}$KIE: Reference | Reference(s) from which $^{13}\varepsilon_{rub}$ is taken from. |
| KIE Assay conditions | Assay conditions at which $^{13}\varepsilon_{rub}$ was measured. All $^{13}\varepsilon_{rub}$ values – with the exception of the *Riftia pachyptila* symbiont rubisco, which was measured at 30°C – were measured at either 25°C or room temperature, so the temperature is omitted. For *Rhodospirillum rubrum*, *Synechococcus elongatus*, and *Spinacia oleracea*, these experiments were conducted by multiple studies in varied reaction conditions that are detailed in Sheet 2. |

Sheet 2 is a compilation of $^{13}\varepsilon_{rub}$ and KIE assay conditions for *Rhodospirillum rubrum*, *Synechococcus elongatus*, and *Spinacia oleracea* only. Values are labeled as follows:

| Column name | Description |
|---|---|
| Reference | Reference from which $^{13}\varepsilon_{rub}$ is taken from. |
| Species | Species of rubisco host. |
| pH | pH at which the KIE assay was performed at. |
| Temperature (°C) | Temperature in degrees Celcius at which the KIE assay was performed. |

| | |
|---|---|
| $[HCO_3^-]$ (mM) | Bicarbonate concentration (mM) at which the KIE assay was performed. |
| RuBP (mM) | RuBP concentration (mM) used in the KIE assay. |
| $[Mg^{2+}]$ (mM) | Magnesium ion concentration (mM) at which the KIE assay was performed. |
| $^{13}\varepsilon_{rub}$ (‰) | Reported mean and standard devation of $^{13}\varepsilon_{rub}$ in units of per mil (‰). All studies except (*27*) report this value as a mean ± s.d., which reports as a median and 95% c.i. in brackets instead. |

**Dataset S2 (separate file).** All data measured in this study. Measured values are labeled as follows:

| Column name | Description |
|---|---|
| Sample_ID | Aliquot number |
| tElapsed_min | Time elapsed since sampling of first aliquot |
| Temp | Temperature (degrees Celcius) |
| Strain | Spinach or *R. rubrum* |
| Area44_first | First eluted Mass 44 Peak Area |
| d13C_avg | Average ($n$=5) $\delta^{13}$C value |
| d13C_sd | Std. dev. ($n$=5) $\delta^{13}$C value |
| Experiment | Experiment name |

Calculated values are labeled as follows:

| Column name | Description |
|---|---|
| volume | Volume calculated from calibration curve |
| volume_error | Propagated error on Volume |
| F | Fraction of DIC pool remaining ($f$) |
| F_err | Propagated error on $f$ |
| R_R0 | $^{13}R_t/^{13}R_i$ |
| R_R0_err | Propagated error on $^{13}R_t/^{13}R_i$ |
| lnF | Natural log of $f$ |
| lnF_err | Error on Natural log of $f$ |
| lnR_R0 | Natural log of $^{13}R_t/^{13}R_i$ |
| lnR_R0_err | Error on Natural log of $^{13}R_t/^{13}R_i$ |
